# The predicted secreted proteome of activated sludge microorganisms indicate distinct nutrient niches

**DOI:** 10.1101/2024.02.27.582363

**Authors:** Kenneth Wasmund, Caitlin Singleton, Morten Kam Dahl Dueholm, Michael Wagner, Per Halkjær Nielsen

## Abstract

In wastewater treatment plants (WWTPs) complex microbial communities process diverse chemical compounds from sewage. Secreted proteins are critical because many are the first to interact with or degrade external (macro)molecules. To better understand microbial functions in WWTPs, we predicted secreted proteomes of WWTP microbiota from more than 1000 high-quality metagenome-assembled genomes (MAGs) from 23 Danish WWTPs with biological nutrient removal. Focus was placed on examining secreted catabolic exoenzymes that target major classes of macromolecules. We demonstrate that Bacteroidota have high potential to digest complex polysaccharides, but also proteins and nucleic acids. Poorly understood activated sludge members of Acidobacteriota and Gemmatimonadota also have high capacities for extracellular polysaccharide digestion. Secreted nucleases are encoded by 61% of MAGs indicating an importance for extracellular DNA and/or RNA digestion in WWTPs. Secreted lipases were the least-common macromolecule-targeting enzymes predicted, encoded mainly by Gammaproteobacteria and Myxococcota. In contrast, diverse taxa encode extracellular peptidases, indicating that proteins are widely used nutrients. Diverse secreted multi-heme cytochromes suggest capabilities for extracellular electron-transfer by various taxa, including some Bacteroidota that encode undescribed cytochromes with >100 heme-binding motifs. Myxococcota have exceptionally large secreted protein complements, probably related to predatory lifestyles and/or complex cell cycles. Many Gammaproteobacteria MAGs (mostly former Betaproteobacteria) encode few or no secreted hydrolases, but many periplasmic substrate-binding proteins and ABC- and TRAP-transporters, suggesting they are mostly sustained by small molecules. Together, this study provides a comprehensive overview of how WWTPs microorganisms interact with the environment, providing new insights into their functioning and niche partitioning.

**Importance:** Wastewater treatment plants are critical biotechnological systems that clean wastewater, allowing the water to reenter the environment and limit eutrophication and pollution. They are also increasingly important for recovery of resources. They function primarily by the activity of microorganisms, which act as a ‘living sponge’, taking-up and transforming nutrients, organic material and pollutants. Despite much research, many microorganisms in WWTPs are uncultivated and poorly characterized, limiting our understanding of their functioning. Here, we analyzed a large collection of high-quality metagenome-assembled genomes from WWTPs for encoded secreted enzymes and proteins, with special emphasis on those used to degrade organic material. This analysis showed highly distinct secreted proteome profiles among different major phylogenetic groups of microorganisms, thereby providing new insights into how different groups function and co-exist in activated sludge. This knowledge will contribute to a better understanding of how to efficiently manage and exploit WWTP microbiomes.

## Introduction

Wastewater treatment plants (WWTPs) play a critical role in removing pollutants and organic matter, and recovering nutrients from wastewater. This is primarily mediated by complex microbiota that degrade, assimilate or transform various organic and inorganic molecules (1–3). Important for this is the organization of microorganisms in WWTPs as multicellular suspended aggregates, known as activated sludge flocs, that facilitate sorption of nutrients to the matrices (4). Organic material can then be biodegraded in the flocs, or eliminated when the flocs are subsequently physically removed (5). Influent waters contain diverse organic material, which provides an array of nutrient sources. The incoming nutrients, as well as molecules produced and recycled *in situ*, play key roles in controlling microbial community compositions by providing nutrient niches for specific guilds that can use specific molecules for growth (6). The degradation of organic matter in activated sludge is often coupled to respiration with electron acceptors such as oxygen or nitrate. This respiration dictates oxygen and nitrate demands of WWTPs, with the latter driving the key process of nitrogen removal through denitrification (7). Further, primary-degraders of organic macromolecules are important for supplying molecules to other key functional guilds such as polyphosphate-accumulating organisms (PAOs) or denitrifiers (8). For example, the fermentative production of acetate provides key PAOs, like *Ca*. Accumulibacter, with their main energy and carbon source (9, 10). Understanding which microorganisms in WWTPs degrade and/or take-up which organic molecules by which mechanisms is therefore critical for understanding how WWTPs function.

Organic matter in WWTPs is largely composed of macromolecules such as proteins (25-35%), carbohydrates (15-25%) and lipids (25-40%) (based on chemical oxygen demand, COD) (6, 11–13). Furthermore, nucleic acids can be abundant, e.g., up to 300 mg of extracellular DNA (eDNA) per g of organic matter in flocs (14). Together, these are generally the most biodegradable macromolecular organics available for microorganisms (15). Additionally, an array of other organic classes occurs in sewage, such as humics/fulvics, steroids, lignins, as well as a large uncharacterised fraction (3). Nevertheless, their turn-over is generally much slower and therefore less influential for WWTP functioning (16–18). Most macromolecules (>600-800 Da MW) need to be digested to smaller components outside of cells, because they are too large to translocate through cell membranes or transporter systems (19). Specific hydrolases and lyases that are secreted or attached to the cell surface are critical for the breakdown of macromolecules. Partially digested molecules are then transported into the periplasm and cytoplasm, where they are further digested to oligomers and monomers, and/or subsequently catabolised or assimilated into new biomolecules (20). Secreted macromolecule-degrading enzymes also promote important ecological interactions, e.g., “cheater” populations may benefit from the degradation products released by primary-degraders without secreting their own hydrolytic enzymes (21). We therefore posited that specifically studying the proteins and enzymes that can be secreted from microorganisms in WWTPs, especially catabolic enzymes, should provide powerful insights into the functioning of microorganisms in WWTPs.

Previous studies have demonstrated the activity of different extracellular hydrolytic enzymes in WWTPs, which are proposed to perform the rate limiting step of organic matter hydrolysis (22). These include peptidases/proteases, phosphatases, esterases/lipases, and carbohydrate-active hydrolases (23–28). Some studies have shown especially high activity of phosphatases, glucosidases and peptidases/proteases (29). Hydrolytic enzyme activities were shown to be persistent over different seasons and conditions (30), including both aerobic and anaerobic phases (31). Extracellular hydrolase activity is mostly associated with flocs and/or extracellular polymeric substances (EPS), suggesting most secreted hydrolases are bound to cells and/or embedded or closely associated with floc matrices (23, 25, 31).

Different hydrolytic activities have also been linked to specific taxa *in situ* (6), mostly using fluorescently-labeled substrates in combination with taxon-specific fluorescent *in situ* hybridisation assays. For example, peptidase activities were linked to cells of phylum TM7 (now Patescibacteria), Chloroflexota and Betaproteobacteria (Pseudomonadota), as well as epiflora of the Saprospiraceaea (Bacteroidota) (32, 33); starch hydrolysis occurred in cells of the Actinobacteriota (Actinomycetota) (34); filamentous Chloroflexi could digest different polysaccharides (35); and lipase activity was linked to members of Mycobacteriaceae (Actinomycetota) (36). While powerful, these studies relied on FISH probes, which limits the observer to the sets of probes applied. Overall, our knowledge on the repertoire of secreted enzymes of functionally important microbes in activated sludge is still very rudimentary and insufficient for understanding nutrient niches in these engineered ecosystems. Genomic-based approaches for predicting secreted hydrolytic activities that are independent of these experimental limitations may prove highly complementary to previous work, potentially offering unique insights into the distributions of enzymes and functional properties of uncultured taxa.

In this study, we aimed to predict and analyze the secreted proteomes encoded among a large collection of high-quality metagenome-assembled genomes (MAGs) from activated sludge from Danish WWTPs that use enhanced nutrient removal technologies (biological N-removal, or N- and P-removal) (37). We investigated two fundamental questions: i) which WWTP microbiota members encode different types of secreted proteins, and ii) which microorganisms have the capacities to drive the primary degradation of organic matter? For the latter, we examined encoded enzymes that may perform catabolic functions for organic macromolecule degradation. Further, we specifically analysed predicted secreted proteins from MAGs representative of abundant populations of WWTPs in Denmark and worldwide (38), as well as members of several functionally relevant populations, such as PAOs, glycogen-accumulating organisms (GAOs), filamentous bacteria, denitrifiers, and nitrifiers. This genome-based information on the secreted proteins can also be directly linked to the Microbial Database for Activated Sludge (MiDAS) (38), because all MAGs analysed contain full-length 16S rRNA genes. Our results provide unique insights into the biology of WWTP microbiota, including a much improved understanding of the capabilities of different taxa to transform organic matter in WWTPs.

## Results and Discussion

### Protein subcellular location profiles predict distinct biology among WWTP microorganisms

To predict secreted proteins and their subcellular locations, we performed subcellular location profiling using PSORTb analysis of all encoded proteins (>4.427 million) from 1083 high-quality MAGs (>90% complete, <5% contamination, and including ribosomal RNA genes) previously recovered from 23 Danish WWTPs (37) (Fig. 1). To focus on proteins from MAGs that represent species-level populations, all results reported below are based on proteins (>2.36 million) from 581 MAGs dereplicated at the species-level (>95% ANI) (37), unless stated otherwise. The MAGs mostly belong to Bacteroidota (35.3%), Gammaproteobacteria (20.1%), Acidobacteriota (5.9%), Actinobacteriota (5.2%), Chloroflexota (5.0%), Myxococcota (5.0%), Alphaproteobacteria (4.8%), and Patescibacteria (4.8%) (Supp. Table 1). The MAGs represent ∼30% of the microbial community members based on relative abundances determined by metagenomic read recruitment (37). Further, most MAGs represent taxa defined as ‘growing’ in the systems (Supp. Table 1), and are thus assumed to be potentially process-critical and not dying-off like many of the incoming influent species (39).

**Figure 1.**
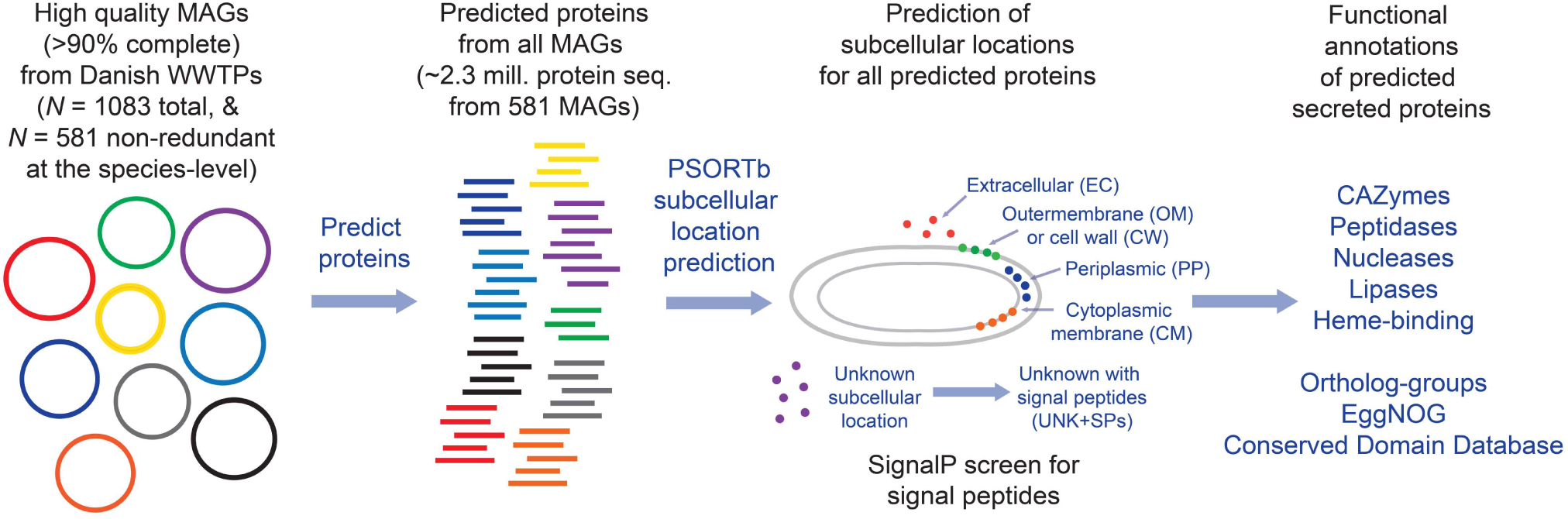
Schematic overview of dataset and analysis pipeline.

**Table 1.** Overview of dataset.

| Protein category | Number | Number with signal peptides |
| --- | --- | --- |
| Total proteins from 581 MAGs | 2,358,707 | N.D. |
| Extracellular | 44,374 | 35,922 (81%)* |
| Outer-membrane | 36,706 | 26,797 (73%) |
| Cell wall | 1,277 | 803 (63%) |
| Periplasmic | 38,428 | 25,559 (67%) |
| Cytoplasmic membrane | 471,235 | N.D. |
| Cytoplasm | 916,404 | N.D. |
| Unknowns | 850,283 | 274,432 (32%) |

\*from three signal peptide detection methods. N.D. Not determined.

The subcellular location profiling predicted 44,374 (1.9%) extracellular, 36,706 (1.6%) outer-membrane, 1,277 cell wall (0.05%), and 38,428 (1.6%) periplasmic proteins (Table 1). Herein, we collectively defined these as “extra-cytoplasmic”. Another 850,283 (36.1%) proteins were assigned “unknown” locations, and 274,432 of these (32.3% of unknown location proteins; 11.6% of all proteins) have signal peptides (here termed “UNK+SP”, i.e., a predicted unknown location and have a signal peptide) indicative of secretion to an extra-cytoplasmic location (Table 1). We analysed UNK+SP proteins separately and in complement to the extra-cytoplasmic proteins. Predicted cytoplasmic-membrane-bound (CM) proteins (471,235; 19.9%) were also analyzed and are briefly described in the Supplementary information.

To additionally gauge the potential of the predicted secreted proteins to be exported from cytoplasms, we independently screened the predicted extracellular proteins for signal peptide sequences for Sec- or TAT-secretion systems (40). Most predicted extracellular proteins (35,921; 81% of extracellular proteins from the 581 non-redundant MAGs) were found to have signal peptides and/or transmembrane features indicative of extra-cytoplasmic locations (Table 1). The prediction of secreted proteins without signal peptides can be explained by the consideration of multiple computational evaluations that PSORTb performs in addition to signal peptide detection, e.g., detection of other motifs and structural signatures, sequence homology to proteins of different subcellular locations, and support vector machine (SVM) analyses of amino acid compositions (41–43). Biological reasons for why extracellular proteins may lack signal peptides include: i) some proteins may be exported via ‘piggy-back’ mechanisms with other proteins/subunits; ii) some secretion systems export proteins with non-canonical signal peptides (e.g., type-9 secretion systems, toxins); and/or, iii) unknown and non-canonical secretion systems and/or signal peptides exist (41, 44).

To investigate the potential of different taxa to secrete different proteins, we determined the numbers of predicted secreted proteins for each of the subcellular locations for each MAG. This aimed to give insights into the propensity of different taxa to secrete proteins to different subcellular locations, which can provide insights about their ecology and functions, e.g., the ability to degrade higher molecular weight organics and/or import degradation products (45). We also explored these numbers of predicted secreted proteins per MAG in relation to the MAG assembly sizes, with the aim to provide an additional perspective on the relative importance of numbers of genes encoding secreted proteins relative to genome sizes among the different taxa.

Among different phyla, MAGs from Myxococcota and Bacteroidota have the most predicted extracellular proteins per MAG, i.e., Myxococcota averaged 202 (SD = 109) per MAG, and Bacteroidota averaged 108 (SD = 48) per MAG (Fig. 2 and Supp. Table 1). Bacteroidota MAGs have high numbers of extracellular proteins for their average MAG sizes (4.5 Mbp, SD = 0.93 Mbp), while many Myxococcota have large MAG sizes averaging 8.2 Mbp (SD = 2.1 Mbp) (Fig. 2, Supp. Fig. 1A and Supp. Table 1). In contrast, MAGs from the Patescibacteria, Elusimicrobiota and Dependentiae generally have the fewest predicted extracellular proteins per MAG, averaging 15 (SD = 9) per MAG, which coincides with relatively small MAG sizes averaging 1.36 Mbp (SD = 0.67 Mbp) (Fig. 2, Supp. Fig. 1A and Supp. Table 1). This fits with the lifestyles of these groups, which are predicted to have relatively simple metabolisms and largely depend on molecules salvaged from other organisms (46–48).

**Figure 2.**
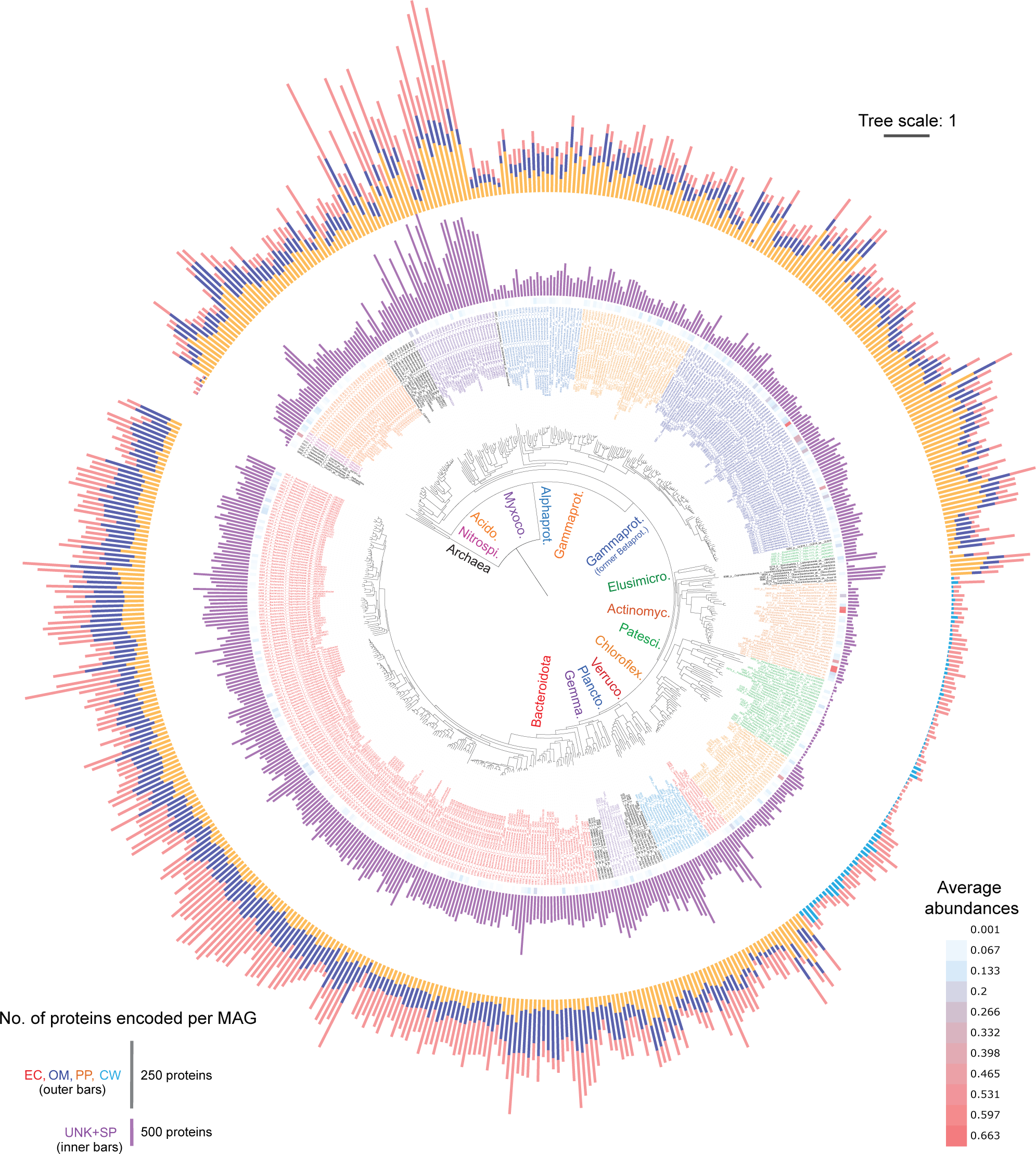
Phylogenomic tree of 581 MAGs from Danish WWTPs with counts of predicted secreted proteins. Outer ring bars (out-to-in) correspond to counts of proteins classified as “extracellular” (red bars, “EC”), “outer-membrane” (blue bars, “OM”), “periplasmic” (orange bars, “PP”), or “cell wall” (teal-blue, “CW”). Scale for counts of proteins are indicated in bottom left legend. Second most inner ring (purple bars, “UNK+SPs”) corresponds to counts of proteins classified as “Unknown with signal peptides”, per MAG. Most inner rign (“Abund.”) with heatmap corresponds to average relative abundances of MAG-populations based on read mapping to MAGs from all metagenomes analysed (values also in Supp. Table 1, colour-scale presented in legend to bottom-right). Leaf labels include the MAG number, followed by taxonomic strings of: phyla (class for Pseudomonadota), family, genus-species, denoted by p, c, f, gs, respectively. Clades of most major phyla are indicated inside the tree with: Nitrospirota; Acidobact. (Acidobacteriota); Myxococc. (Myxococcota); Alphaprot. (Alphaproteobacteria); Gammaprot. (Gammaproteobacteria); Betaprot. (Betaproteobacteria); Elusimicro. (Elusimicrobiota); Actinomyc. (Actinomycetota); Patesci. (Patescibacteria); Chloroflex. (Chloroflexota), Verruco. (Verrucomicrobiota); Plancto. (Planctomycetota); Gemma. (Gemmatimonadota); Bacteroidota. GTDB species names are only presented if named, i.e., GTDB number codes were removed. The tree is based on a concatenated alignment of protein sequences derived from single copy marker genes obtained from CheckM analysis of MAGs. Scale bar represents 100% sequence divergence.

MAGs of the Bacteroidota (including class Ignavibacteria) and several Acidobacteriota MAGs, have “especially high numbers” (defined throughout as those with >2SD above the mean of counts per MAG, among all non-redundant MAGs) of predicted outer-membrane proteins per MAG (ranging 139-174 per MAG) (Fig. 2, Supp. Fig. 1C and Supp. Table 1). Chloroflexota had the highest numbers of predicted cell wall-bound proteins among organisms with Gram-positive cell walls (up to 65) (Fig. 2, Supp. Fig. 1C and Supp. Table 1). Groups encoding high numbers of predicted periplasmic proteins per MAG included Gammaproteobacteria (mostly from Burkholderiales, i.e., former Betaproteobacteria), Myxococcota, and an Acidobacteriota MAG (ranging 170-283 per MAG) (Fig. 2, Supp. Fig. 1D and Supp. Table 1). For predicted CM proteins, Chloroflexota MAGs stood-out for having especially high numbers, averaging 1725 per MAG (SD = 524) (Supp. Table 1, Supp. Fig. 1E and Supp. Fig. 2).

Numbers of UNK+SP proteins per MAG correlated strongly with total numbers of extra-cytoplasmic proteins per MAG (Pearson correlation, *R* = 0.841, *p* < 0.001) (Supp. Fig. 3). This supports the strong trends in the propensity of different taxa with capabilities to secrete varying types of proteins. MAGs of Myxococcota and Bacteroidota had the most, averaging 1186 (SD = 431) and 584 (SD = 133) UNK+SP proteins per MAG, respectively (Fig. 2, Supp. Fig. 1E and Supp. Table 1). MAGs from the Patescibacteria generally have the fewest predicted UNK+SP proteins per MAG, averaging only 44 (SD = 15). Notably, MAGs from several phyla had many UNK+SP proteins that had few PSORTb-predicted extra-cytoplasmic proteins, i.e., the Planctomycetota, Gemmatimonadota, Acidobacteriota and Verrucomicrobiota, among others (Fig. 2 and Supp. Table 1). This suggests they have many proteins that are not similar to proteins that have been proven to be secreted, and were therefore given “unknown” locations by PSORTb, although they are likely secreted. This implies that these taxa are enriched in novel secreted proteins, which likely impart undescribed functions.

### Secreted carbohydrate active enzymes are most prevalent among Bacteroidota

Carbohydrates are abundant in WWTPs, accounting for 18% of the COD in influent wastewater (49), and of which up to 85% is high molecular weight (50). They are derived from the supply of fresh sewage material (49), or from *in situ* produced extracellular polymeric substances (EPS), and/or cellular components of biomass (51). They are therefore major nutrient sources for WWTP microbiomes, with high removal rates of up to 85% of carbohydrates from wastewater indicating they are readily biodegraded (52). We therefore identified carbohydrate active enzymes (CAZy) that are predicted to be secreted and could help microorganisms to degrade and use polysaccharides, i.e., glycoside hydrolases, proteins with carbohydrate-binding modules, carbohydrate esterases, and polysaccharide lyases (Supp. Table 5).

Overall, Bacteroidota encode the most extracellular CAZy proteins per MAG, e.g., they represent 20 of the top 26 MAGs when ranked by numbers of extracellular CAZy enzymes (i.e., those with >2SD above the mean; ≥8 extracellular CAZy per MAG) (Supp. Table 5). Additionally, two Verrucomicrobiota MAGs, two Fibrobacterota MAGs, and a single *Cellvibrio* MAG were also among MAGs encoding high numbers of extracellular CAZy enzymes (>2SD above the mean). These taxa likely have specialized capabilities to degrade high-molecular weight polysaccharides.

MAGs with multiple (≥2) predicted outer-membrane-bound CAZy mostly belonged to Bacteroidota (*N* = 58), as well as Gemmatimonadota (*N* = 7), Verrucomicrobiota (*N* = 2), Acidobacteriota (*N* = 2), and single MAGs of Fibrobacterota, Gammaproteobacteria, Alphaproteobacteria and Planctomycetota (Supp. Table 5). MAGs of the Ignavibacteria (Bacteroidota) encode the most, with up to 9 outer-membrane CAZy per MAG. Among the Gram-positive lineages, cell wall-associated CAZy were encoded by various Chloroflexota (Anaerolineae) (*N* = 18), and several Actinomycetota (*N* = 6) and Patescibacteria (*N* = 4) (Supp. Table 5). Similarly, MAGs of Bacteroidota (*N* = 3), Verrucomicrobiota (*N* = 2), a Gemmatimonadota MAG and a Acidobacteriota MAG were among MAGs encoding especially high periplasmic CAZy (those with >2SD above mean) (Supp. Table 5).

Among the different classes of extra-cytoplasmic CAZy, glycoside hydrolases were most widespread among MAGs (*N* = 387), followed by carbohydrate-binding modules (*N* = 155), carbohydrate esterases (*N* = 154) and polysaccharide lyases (*N* = 47) (Supp. Table 5). Similarly, among UNK+SP proteins, glycoside hydrolases were the most widespread among MAGs (*N* = 427), while polysaccharide lyases were the least common (*N* = 115) (Supp. Table 5). This suggests fewer taxa have the capacity to use polysaccharides with complex structures and/or modifications that require enzymes like carbohydrate esterases and polysaccharide lyases.

In total, 17 MAGs encode all aforementioned extra-cytoplasmic CAZy types, and belonged to the Bacteroidota (mostly genus *Haliscomenobacter*), a Fibrobacterota MAG, a Verrucomicrobiota MAG and a Gammaproteobacteria MAG (Supp. Table 5). An additional 32 MAGs encode all four major CAZy types among UNK+SP proteins, and mostly belong to diverse Bacteroidota (*N* = 19), as well as Myxococota MAGs (*N* = 5), several Gammaproteobacteria, Fibrobacterota and Acidobacteriota MAGs, and single MAGs of Krumholzibacteriota and Planctomycetota (Supp. Table 5). These taxa likely have capacities to degrade structurally complex polysaccharides that require multiple types of CAZy.

Together, the results show that diverse and abundant Bacteroidota have high capacity to contribute to digesting diverse extracellular polysaccharides, which indicates they probably make important contributions to the process *in situ*. This is in line with the known ability of many Bacteroidota as polysaccharide-degrading specialists in marine and mammalian gut systems (53). Several members of the Chloroflexota are also abundant, probable extracellular polysaccharide-degraders, which supports previous work in WWTPs (35). Other taxa with probable extracellular polysaccharide-degrading capabilities from the Verrucomicrobiota, Planctomycetota, Fibrobacterota, Acidobacteriota and *Cellvibrio* are also known for their polysaccharide-degrading capabilities in other environments (54–57), while taxa of Gemmatimonadota that have high numbers of secreted CAZymes are poorly understood with regards to carbohydrate use.

### Peptidases are the most prevalent secreted hydrolytic enzymes encoded

Proteins are among the most abundant and labile nutrients available for microorganisms in WWTPs (49). To identify secreted peptidases and/or proteases (herein ‘peptidases’) with probable “nutrient-acquiring” functions, we identified predicted secreted peptidases and took conservative steps to exclude peptidases likely associated with biosynthetic or house-keeping functions, i.e., we mapped peptidases to different functional categories of clusters of orthologous groups (COG) and excluded those mapping to biosynthetic or house-keeping categories (see Materials and Methods). From this, we identified 572 predicted extracellular peptidases among 291 MAGs from diverse taxonomic groups, i.e., 21 of 33 phyla and classes of Pseudomonadota (Proteobacteria) (Supp. Table 1). Many (55%) belonged to MEROPS peptidase family M4, which includes homologs to bacillolysin/thermolysin-type peptidases that are known as secreted “nutritional” peptidases (58). MAGs from the Bacteroidota (*N* = 25), Acidobacteriota (*N* = 4), Gammaproteobacteria (*N* = 3), Myxococcota (*N* = 3) and a Planctomycetota MAG encoded high numbers of extracellular peptidases per MAG (>2SD above the mean; ≥4 per MAG). Some of these taxa (Bacteroidota, Acidobacteroidota and Planctomycetota) were previously identified as enriched with genes for secreted peptidases among various environments (59). Among MAGs that encode numerous extracellular peptidases, members of the *Thermomonas* (Gammaproteobacteria) are known protein-degraders (60). This supports the notion that the predicted secreted peptidases they encode, and related types from other taxa, could be used for extracellular protein digestion.

We also used the same classification strategy for peptidases with predicted outer-membrane and cell wall locations. Fewer peptidases with predicted outer-membrane/cell wall locations were found (*N* = 134), mostly in the same taxa (*N* = 107) that encode extracellular peptidases (Supp. Table 1). Among predicted periplasmic and UNK+SP peptidases, most were related to peptidases with probable house-keeping functions and were therefore not analysed further.

Overall, these results indicate widespread potential to secrete peptidases by diverse taxa in WWTPs. Extracellular peptidases were the most common type of predicted extracellular catabolic enzymes targeting any of the major macromolecule classes in this study. This is in line with previous enzymatic assays in WWTPs that showed protease activity was the highest among macromolecule-degrading activities tested (23, 27). We nevertheless wish to point-out that differentiating secreted peptidases with nutrient-acquiring functions versus biosynthetic or house-keeping functions, should be treated with caution.

### Secreted lipases indicate capacity to use lipids among a select subset of taxa

Lipids and fats are abundant in influent water of WWTPs making up to 40% of COD (49), and secreted lipases are needed to initiate their breakdown. Predicted extracellular lipases were identified in 153 MAGs (Supp. Table 1). MAGs from the Myxococcota (*N* = 10), Gammaproteobacteria (*N* = 9), Bacteroidota (*N* = 3), two Alphaproteobacteria (*N* = 2) and a Bdellovibrionota MAG encoded multiple (≥2 per MAG) copies of extracellular lipases with signal peptides and/or transmembrane features, with up to 5 encoded by *Rhodoferax* MAG 0761. Predicted outer-membrane lipases were restricted to 63 MAGs, and were most common among Gammaproteobacteria MAGs (*N* = 40), although MAGs from a few other groups including family PHOS-HE28 of Bacteroidota encoded outer-membrane lipases, too. Important to note, is that the functional roles of outer-membrane-bound lipases are not completely understood, with some studies suggesting they could be involved in cell-membrane repair (61). Only eight MAGs had predicted cell wall-bound lipases, with six MAGs from Actinomycetota families and two MAGs of Caldilineaceae (Chloroflexota). An additional 158 MAGs spanning diverse taxa had predicted lipases among UNK+SP proteins, with 109 of these MAGs not having any predicted extracellular lipases (Supp. Table 1). The Bdellovibrionota MAG 0471 encoded the most, with 8 lipases among UNK+SP proteins. There were 44 MAGs with multiple (≥2 per MAG) lipases among UNK+SP proteins, most being Gammaproteobacteria (*N* = 14), Myxococcota (*N* = 11), and Actinomycetota (*N* = 5).

Many of the MAGs encoding predicted secreted lipases are related to taxa known to have lipase activity, thereby supporting the functional predictions made here. For example, MAGs from the known lipolytic gammaproteobacterial genera *Agitococcus* (MAG 1031) (62) and *Rhodoferax* (MAG 0761) (63) encoded 3 and 5 extracellular lipases with SPs, respectively. *Ca.* Microthrix (Actinomycetota) have been shown to be specialized long-chain fatty acid-degraders (i.e., oleic acid) in situ and in vitro in activated sludge (64, 65), and all MAGs of this genus encoded predicted extracellular and/or UNK+SP lipases. We hypothesize secreted lipases among members of the Myxococcota and Bdellovibrionota may be involved in digesting the cell wall lipids of their prey (66, 67), because these taxa often exhibit predatory lifestyles (67–69). The lower number of MAGs with secreted lipases compared to hydrolases for the other major classes of macromolecules suggests a more specialized range of taxa have capacity for degradation of lipids, than for the other classes of macromolecules.

### Secreted nuclease genes are common suggesting important functional roles

Extracellular nucleic acids may act as sources of nutrient or nucleic acid building blocks, and/or may play structural roles within biofilm-like flocs in WWTPs. Nevertheless, nothing is known about nucleic acid-degrading taxa in WWTPs. Overall, we identified diverse MAGs from various phyla (*N* = 351 MAGs; 61% of MAGs) that encode predicted secreted nucleases, i.e., DNases and/or RNases (Supp. Table 1). Different types of extracellular nucleases (excluding specific RNases; see below) were generally encoded by different phyla, e.g., endonuclease-type by Bacteroidota; NUC-type nucleases by Bacteroidota and Acidobacteiota; SNc-type nucleases by Gram-positive Actinomycetota, Chloroflexota and Patescibacteria; and HNHc-type nucleases by diverse taxa (Supp. Table 6). MAGs of the Bacteroidota, Acidobacteriota and Chloroflexota were notable because they comprised 44 of the 51 MAGs that encoded multiple (≥2) predicted extracellular nucleases. Previous work in marine sediments showed that bacteria with multiple copies of genes for extracellular nucleases were active DNA-degraders within experimental microcosms (70). Interestingly, 15 of the 28 Patescibacteria MAGs encoded extracellular nucleases. This is noteworthy because all Patescibacteria MAGs had few other predicted secreted proteins. Because Patescibacteria typically lack biosynthetic capabilities (71, 72), we hypothesize they use them to help salvage nucleobases for incorporation into new nucleic acids.

In total 123 MAGs encoded probable secreted nucleases among UNK+SPs proteins (Supp. Table 1). Seventeen of these had multiple (≥2) nucleases among UNK+SP proteins, including several Bacteroidota and Myxococcota MAGs, and single MAGs of Krumholzibacteriota, Eisenbacteria, Planctomycetota and Gammaproteobacteria (Supp. Table 1). Few nucleases were predicted to be outer-membrane-bound (*N* = 10), and were mainly present among Bacteroidota. Predicted cell wall-bound nucleases were common among MAGs of Gram-positives, with 50% and 59% of Actinomycetota and Chloroflexota MAGs encoding them, respectively. Predicted periplasmic nucleotidases were widespread among MAGs (*N* = 240), being common among MAGs of Bacteroidota (*N* = 105). Many Gammaproteobacteria MAGs (*N* = 36) also encoded predicted periplasmic nucleotidases, but lacked other extracellular or outer-membrane nucleases. This suggests they are equipped to use free nucleotides but not to degrade polymeric nucleic acids.

Predicted secreted RNases were more restricted, i.e., we identified 21 MAGs with extracellular RNases, and 69 MAGs with RNases among UNK+SP proteins (Supp. Table 1). Extracellular RNases were encoded in various Actinomycetota (*N* = 11), several Chloroflexota (*N* = 4) and Patescibacteria MAGs (*N* = 3), and single Bacteroidota and Firmicutes MAGs. Many of the MAGs with RNases among UNK+SP proteins were members of the Burkholderiales (*N* = 58) (Supp. Table 1). No RNases were identified for proteins among any of the other predicted extra-cytoplasmic compartments.

Together, these results suggest extracellular nuclease activity might be an important yet underappreciated aspect of WWTP microorganisms. Nucleic acids could be supplied by the mass immigration of microorganisms into activated sludge that then die-off (39), or from in situ production for floc structures, or from in situ derived necromass. The ability to degrade nucleic acids could be important for processes such as: i) acquiring molecules such as bases or ribose for catabolism, ii) enabling salvage of nucleobases, iii) acquisition of phosphorus, and/or iv) regulating the structures of activated sludge flocs that contain extracellular DNA, if similar to these functions in other biofilms (73).

### Heme-binding proteins predict major differences in redox properties of WWTP microorganisms

Secreted heme-binding proteins including cytochromes can mediate diverse electron transfer reactions and indicate capabilities to perform redox reactions and/or tolerate changing redox conditions. We therefore searched for the common canonical heme-binding motif (CxxCH) (74) among predicted secreted proteins, and identified those with similarities to cytochromes. Further, we specifically identified proteins with multi-heme-binding sites (≥4) with predicted extracellular/outer-membrane/cell wall locations, because they often mediate extracellular electron transfer (75). We identified 259 multi-heme proteins among 149 MAGs, with MAGs encoding numerous multi-heme proteins (>2SD above the mean; ≥3 per MAG) belonging to the Bacteroidota (*N* = 11), Myxococcota (*N* = 4), Gammaproteobacteria (*N* = 3), Planctomycetota (*N* = 2), phylum AABM5-125-24 (*N* = 2), as well as single MAGs of the Verrucomicrobiota, Acidobacteriota, Desulfobacterota and “JADJOY01” (Fig. 3, Supp. Table 7). Many of these MAGs are from groups known to encode extracellular cytochromes for mediating extracellular electron transfer, such as *Anaeromyxobacter* (Myxococcota), Geobacterales (Desulfuromonadota) (76) and *Geothrix* (Acidobacteriota) (77). Multi-heme cytochromes were also previously reported in the same *Geothrix*-related MAGs (78), although sub-cellular locations were not predicted. Recent experimental work showed increases of *Geothrix* and Ignavibacteria spp. (Bacteroidota) in WWTPs when dosed with Fe(III) under anaerobic conditions (79), suggesting they used Fe(III) as an electron acceptor for growth. We therefore suggest the extra-cytoplasmic cytochromes we identified could facilitate such reactions.

**Figure 3.**
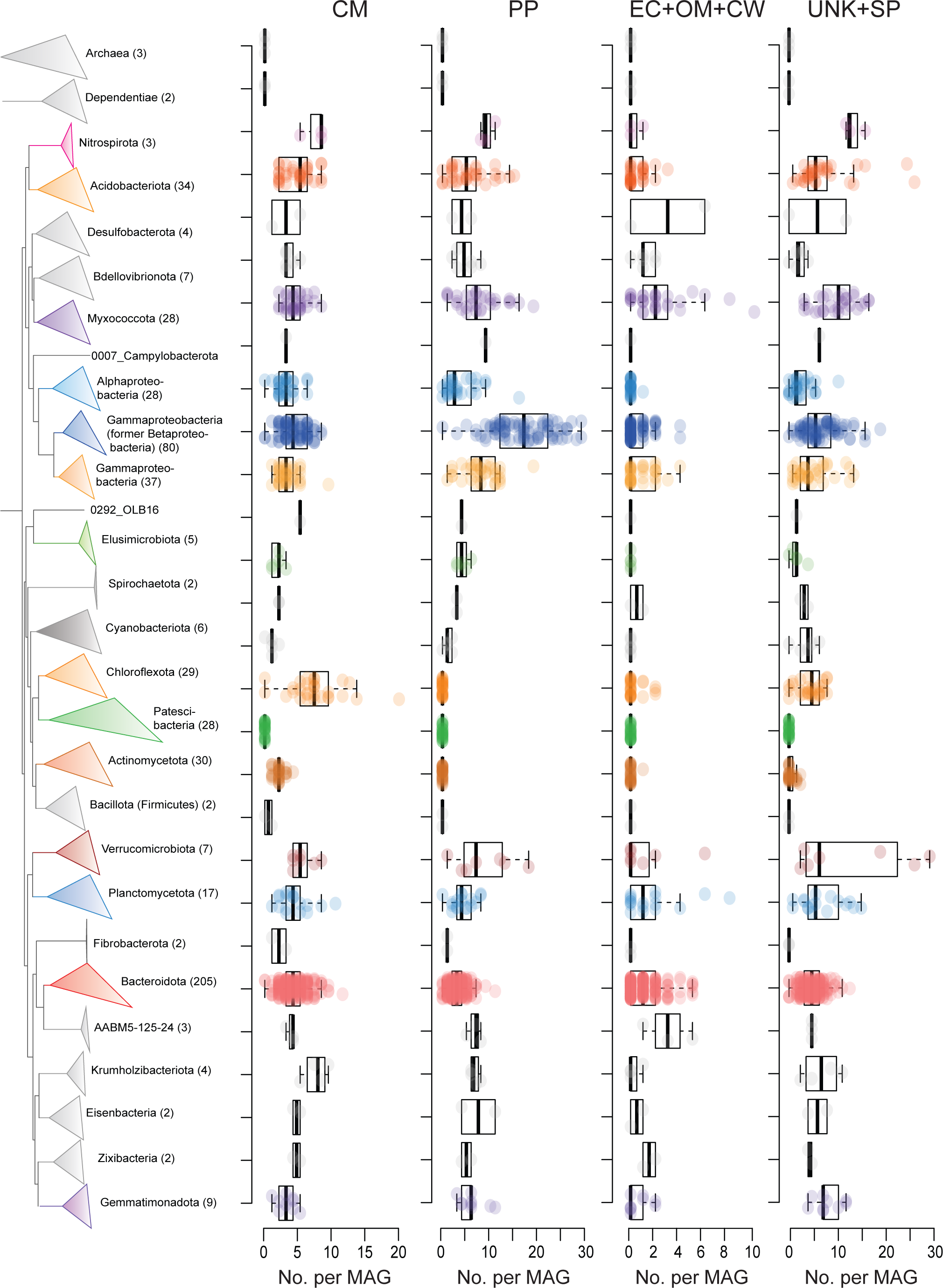
Phylogenomic tree of MAGs from Danish WWTPs with counts of heme-binding proteins for MAGs the major groups. Numbers in parenthesis of taxa names indicate the number of MAGs in each group. Counts are presented for proteins predicted to be present in the cytoplasmic-membrane (CM), periplasmic (PP), extracellular+outer-membrane+cell wall (EC+OM+CW), or unknown location with signal peptide (UNK+SP). For EC+OM+CW and UNK+SP heme-binding proteins, counts of predicted multi-heme proteins (≥4 heme-binding sites) are presented. For PP and CM heme-binding proteins, counts of proteins with ≥1 heme-binding sites are presented. Boxplots show the summary statistics with boxes indicating interquartile ranges (IQR), whiskers indicate range of values within 1.5 × IQR, and horizontal lines show medians. The branches linking Archaea and Dependentiae to the tree are not shown, where the Dependentiae branch with the rest of the Bacteria.

Analysis of UNK+SP proteins identified 708 multi-heme-binding proteins among 295 MAGs (Fig. 3, Supp. Table 7). Groups with many (>2SD above the mean) multi-heme-binding proteins per MAG belonged to Myxococcota, Acidobacteriota, Planctomycetota, Chloroflexota and Krumholzibacteriota, among a few others. The arrays of multi-heme binding proteins highlight taxa in WWTPs that could potentially mediate electron exchange between insoluble molecules such as insoluble metals, humic-like organics, or directly between other cells. These findings also indicate previously unrecognized capacity for extracellular electron exchange among various taxa in WWTPs, especially among Bacteroidota.

Among predicted secreted multi-heme-binding proteins, we identified many proteins with high numbers of heme-binding sites per protein among MAGs (Supp. Table 8). They are especially prevalent among uncharacterised genera of the *Saprospiraceae* (Bacteroidota), e.g., they comprised 90% of the MAGs among the 50 proteins with the most heme-binding sites. A *Paludibaculum* MAG (Acidobacteriota) contained the most for a single protein, with 112 heme-binding motifs. We speculate they may play roles in extracellular electron transfer or electron storage “capacitor-like” functions (80).

Among predicted periplasmic proteins with heme-binding sites, we identified diverse *c*-type cytochromes encoded in especially high numbers (>2SD above the mean; ≥18 per MAG) among MAGs of the Burkholdariales (*N* = 34) (Gammaproteobacteria), and single MAGs of Chromatiales (Gammaproteobacteria), Myxococcota and Verrucomicrobiota (Supp. Table 7). High numbers and diversity of cytochromes likely impart physiological flexibility through redox flexibility (81, 82). Overall, these results suggest major differences among different phylogenetic clades in their ability for cytochrome-mediated respiratory flexibility and/or abilities to tolerate changes in redox conditions in WWTPs. These properties likely manifest in differences in metabolic activity and/or ecological success under the fluctuating redox conditions of activated sludge, which undergo drastic switches between anoxic and oxic conditions.

### Myxococcota MAGs encode especially large complements of secreted proteins

Myxococcota MAGs encode the highest numbers of predicted extracellular proteins and UNK+SPs among all MAGs (Fig. 2 and Supp. Table 1), and therefore we aimed to explore the complements of their predicted extracellular proteins (apart from hydrolytic enzymes). Although it was beyond the scope of this study to analyze all predicted extracellular proteins from Myxococcota in detail, our analyses revealed: i) an expansive array of protein sequence diversity with little similarly to proteins with known functions; ii) various unusual proteins that are seemingly enriched among Myxococcota and few other bacterial phyla, but also present in eukaryotes, i.e., proteins with Stigma1 domains often found in proteins from fungi and plants; iii) many secreted proteins with adhesion properties that might be important for their functioning. Further details are described in the Supplementary information.

### Key features of predicted secreted proteomes of the most abundant taxa and key functional groups

Finally, we specifically analyzed predicted extra-cytoplasmic and UNK+SP proteins from MAGs (*N* = 63) that represent abundant, as well as functionally relevant taxa (defined above) (Fig. 4), i.e., taxa likely relevant to nutrient removal processes such as PAOs, GAOs and nitrogen cycling bacteria like nitrifiers and denitrifiers (see Methods). First, we performed ortholog-group (OG) analysis of extra-cytoplasmic proteins and UNK+SP proteins, separately, to identify highly-represented types of secreted proteins encoded among these taxa (Supp. Table 1). From these, we explored the functions of proteins from the top 100 OGs of these MAGs, i.e., OGs were ranked by sums of counts of proteins from MAGs, among each OG (Supp. Table 2 and Supp. Table 3). Hierarchical clustering of OGs and MAGs revealed clear phylogenetic clustering of MAGs based on OG contents of extra-cytoplasmic proteins (Fig. 5). This highlights that phylogenetically related microbes that are abundant and/or share similar process functions in activated sludge encode similar types of secreted proteins.

**Figure 4.**
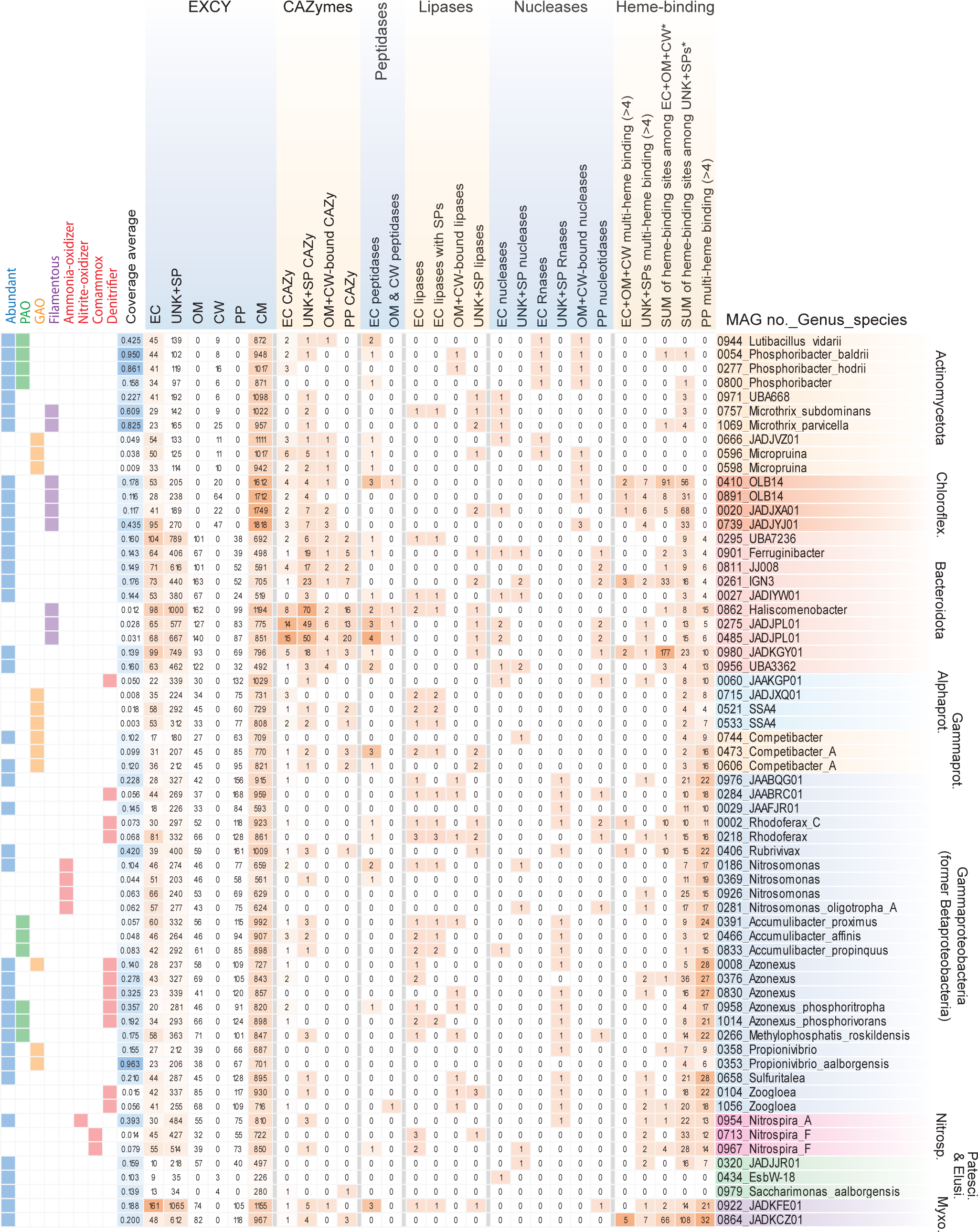
Heatmap of the counts of proteins of different secreted compartments, and for CAZymes, peptidases, lipases, nucleases and heme-binding proteins from abundant and functionally relevant taxa. Numbers of proteins per MAG are indicated in each cell. Colour scales were set for each of extracytoplasmic (EXCY) columns separately, while the colour scale for all other catabolic proteins and heme-binding are set separately. Clades of most major phyla are indicated with abbreviations being: Chloroflex. (Chloroflexota); Alphaprot. (Alphaproteobacteria); Gammaprot. (Gammaproteobacteria); Patesci. (Patescibacteria); Elusi. (Elusimicrobiota); Myxoco. (Myxococcota).

**Figure 5.**
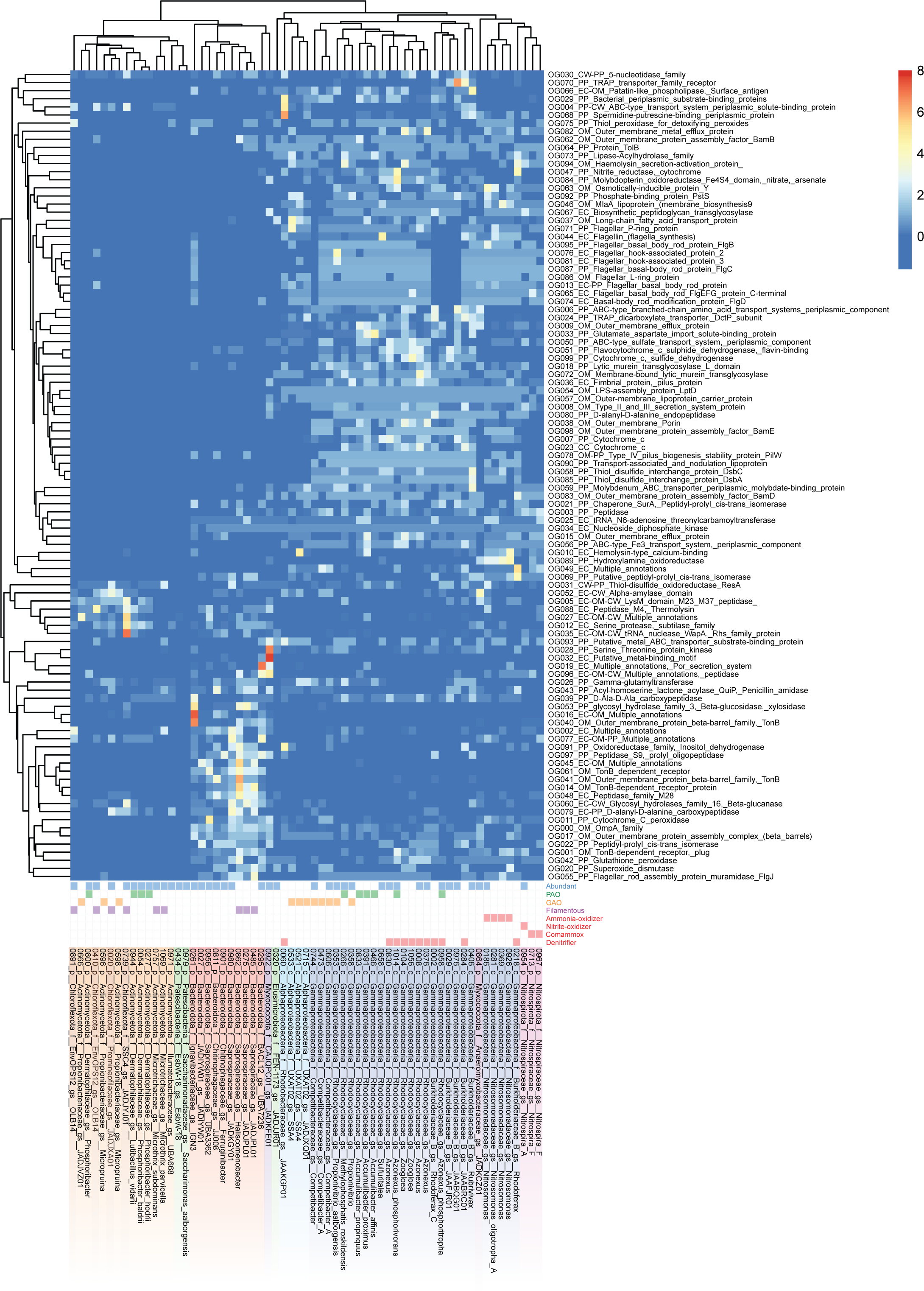
Heatmap and cluster analysis of numbers of encoded proteins from orthogroup analysis of secreted proteins from abundant and functionally relevant taxa, i.e., how many proteins of each OG were encoded per MAG. Rows are centered; unit variance scaling is applied to rows. Both rows of OGs and columns of MAGs are clustered using correlation distance and average linkage using ClustVis (108). The scale for the heatmap colours are indicated in the legend, where the scale maximum of 8 was used to enhance visualisation and differentiation of lower values <8. Note that some values were therefore >8, and raw values of OG counts are available in Supp. Table 2. MAG labels include the MAG number, followed by taxonomic strings of: phyla (class for Pseudomonadota), family, genus-species, denoted by p, c, f, gs, respectively.

The highest represented OGs related to proteins involved in nutrient catabolism and import (excluding probable biosynthetic or housekeeping functions) included proteins of TonB transporter outer-membrane barrels and receptors that were abundant among Bacteroidota and Acidobacteriota MAGs, while components of ABC/TRAP proteins were common among Gammaproteobacteria MAGs (Fig. 5 and Supp. Table 2). This included most denitrifiers, which are mostly from the Gammaproteobacteria. Among OGs of UNK+SPs proteins, components of ABC-transporter proteins were also identified among Gram-positive Actinomycetota and Chloroflexota MAGs (Supp. Fig. 4 and Supp. Table 3). TonB receptors are often associated with polysaccharide import systems and/or transport of larger organic molecules including vitamins and siderophores (83), while ABC- and TRAP-transporters are generally thought to be used for import of smaller low-molecular weight organics (84–86). Thus, this suggests distinct organic substrate preferences among these major phylogenetic groups in activated sludge. Additional findings regarding potential catabolic enzymes and proteins potentially involved in interspecies competition are described in the Supplementary information. OG analysis of predicted cytoplasmic membrane proteins were briefly explored and are described in the Supplementary information (Supp. Fig. 5 and Supp. Table 4).

Next, we specifically examined secreted macromolecule-degrading enzymes predicted from the abundant and functionally relevant taxa (Fig. 4). We revealed that different PAOs encode secreted enzymes that may facilitate contrasting ecological strategies. For instance, *Ca*. Phosphoribacter (MAGs 0054 and 0277) and *Ca*. Lutibacillus (MAG 0944) (all formerly “*Tetrasphaera”*) (87), encode suits of secreted catabolic enzymes for different macromolecules, i.e., peptidases, RNases and CAZymes. They are differentiated from other PAOs like *Ca*. Accumulibacter and some *Azonexus* (*Ca*. Dechloromonas) that instead have predicted secreted lipases and very few CAZymes. This may be important for niche differentiation among PAOs. Key nitrogen cycling organisms such as ‘nitrifiers’ (ammonia- and/or nitrite-oxidizers, including complete ammonia-oxidizers, i.e., ‘comammox’), encode very few secreted proteins and/or catabolic enzymes, which is in line with their specialized chemolithotrophic lifestyles that would not require investment in secreted hydrolases.

Many abundant Gammaproteobacteria MAGs (18 of 28) encoded predicted RNases with signal peptides (UNK+SPs), and some had lipases. In contrast, many of the abundant filamentous bacteria in activated sludge (i.e., Chloroflexota, Actinomycetota and Bacteroidota) have the capacity to be primary-degraders of organic macromolecules, whereby most encode numerous secreted catabolic enzymes for macromolecules.

Among the abundant MAGs, two abundant Myxococcota MAGs had contrasting features. Anaeromyxobacteraceae MAG 0864 had the second most predicted extracellular cytochromes of any MAG (*N* = 8), and also many predicted periplasmic cytochromes (*N* = 32) (Fig. 5). This indicates high redox flexibility, as discussed previously for a cultured relative *Anaeromyxobacter dehalogenans* (91). The other MAG 0922 of an uncultured class GTDB UBA796 has the largest array of predicted extracellular proteins (*N* = 161) among the most abundant/core organisms analyzed here (Supp. Table 1). It has extensive extracellular hydrolytic potential for digesting macromolecules, encoding three predicted extracellular peptidases, extracellular and UNK+SP CAZymes, as well as potential lipases and nucleases among UNK+SP proteins.

Additional findings from manual inspections of secreted protein annotations of abundant taxa and key functional groups are detailed in the Supplementary information.

## Conclusions

This study shows that predicted secreted proteins encoded by genomes of activated sludge microorganisms are highly distinct across different taxonomic groups, which indicates unique and contrasting ecological strategies, as well as potentially unique niche spaces (Fig. 6). We find strong evidence for the potential to digest extracellular macromolecules by key filamentous bacteria of Actinomycetota and Chloroflexota, many Bacteroidota, as well as key PAOs of *Ca*. Phosphoribacter and *Ca*. Lutibacillus (former *Tetrasphaera*). These taxa are therefore likely functioning as primary-hydrolysers in the microbial food webs of WWTPs. In contrast, most Gammaproteobacteria (mostly Burkholderiales, former Betaproteobacteria), many of which are abundant and/or functionally relevant populations, have limited capacity for extracellular hydrolysis of macromolecules, but seem adapted to utilize smaller and simple organics. Our analyses highlight Bacteroidota as key polysaccharide-degraders, but also groups that are poorly understood in activated sludge including Gemmatimonadota, Myxococcota and Acidobacteriota. We find that peptidases are the most taxonomically widespread secreted hydrolytic enzymes, while secreted lipases are the most restricted. We also show that secreted nucleases are encoded by diverse bacteria, suggesting important functions. Finally, our results provide a catalog of the secretion potential of all the MAGs investigated that can be linked to the MiDAS database (Supp. Table 1) representing the majority of all abundant genera in WWTPs worldwide (38). Overall, this study reveals new perspectives into the functional potential of microorganisms in WWTPs and their potential to interact with the external environment. Future studies are needed to experimentally confirm the predictions made.

**Figure 6.**
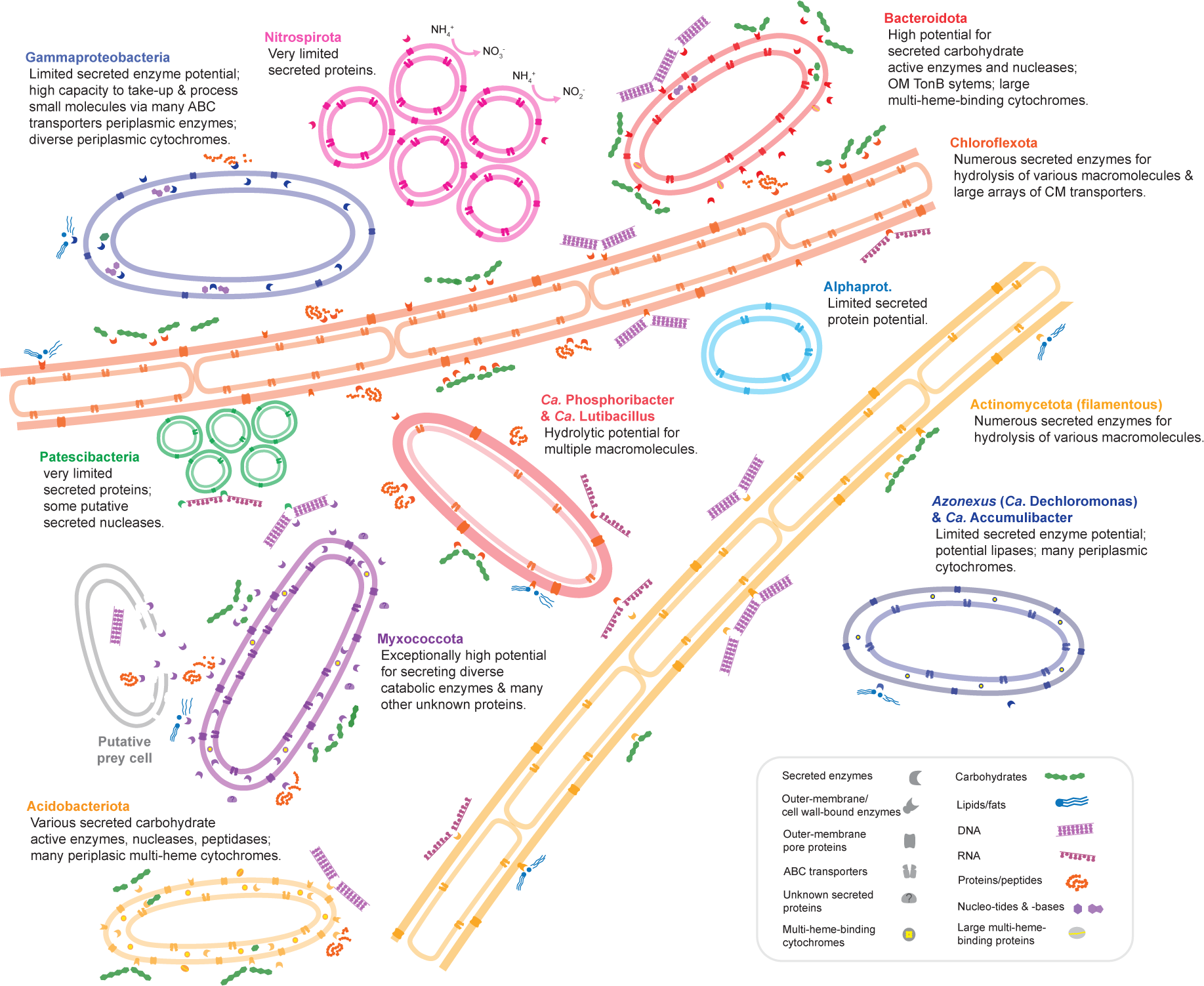
Schematic depiction of key and general findings for abundant and key functional taxa.

## Materials and Methods

### Subcellular location profiling of proteins

The 1083 MAGs analyzed in this study were previously generated from WWTPs across Denmark (37). The MAGs were given code numbers from 0001 to 1083 (Supp. Table 1). The taxonomies of the MAGs used in this study were obtained using GTDB-tk (v2.3.0) (88) with database release 214 (89). A phylogenomic tree of MAGs was generated using the maximum-likelihood algorithm using IQ-TREE web-server with automatic substitution model and ultra-fast bootstrapping (1000×) (90). The tree was based on the alignment of concatenated protein sequences derived from single copy marker genes retrieved from CheckM (91). The tree was curated with iTol (92). In this study, when describing taxa at phylum level, the classes of Pseudomonadota (syn. Proteobacteria) were described in place of the phylum, in order to better explore the specific properties of the classes of this diverse group, and to provide readers information regarding traditionally used taxonomies. The MAGs were subject to initial automatic protein calling and annotation using Prokka (v1.14.5) (93) with default settings. The subcellular locations of all encoded proteins were then predicted using PSORTb 3.0 (v3.0.6) (41) with the option for cell wall/membrane types set for each MAG (Supp. Table 9). Cell wall types were used according to PSORTb pre-computed profiles for known phyla, while literature searches were done to set the cell wall type option for newly described and uncultured taxa when such information was available. We chose sub-cellular location prediction as the strategy to predict secreted locations rather than only predicting secreted proteins with secretory signal peptides (SEC or TAT), because sub-cellular location prediction gives information regarding the probable final locations of the proteins, and many secreted proteins lack signal peptides (41). PSORTb was chosen because it enables high-throughput analysis, is accurate, and takes into account different cell wall types (41). Protein sequences given “Unknown” locations by PSORTb were also collected and subjected to SignalP (v 5.0b) (40) analysis using the options for the same cell wall types as per for MAGs subject to PSORTb (described above). This was done to predict additional, probable secreted proteins containing signal peptides (SEC or TAT). SignalP was chosen because it can facilitate high-throughput analysis. For additional and specific analysis of signal peptides among predicted “extracellular” proteins, we performed SignalP analysis as described above, as well as with the PRED-TAT server (94) using “original model”, and using Phobius (v1.01) (95) using default settings.

### General annotations of proteins

For subsets of proteins specified, additional and complementary functional annotations of proteins were obtained using eggNOG-mapper (v2.0.0) (Cantalapiedra et al. 2021) using default settings (minimum hit e-value 0.001, minimum hit bit-score 60, minimum % of identity 40, minimum % of query coverage 20) with the “-m diamond” option. Where specified, protein sequences were screened for conserved domains using the Conserved Domain search tool (96) against the Conserved Domain Database (CDD) (97) with default settings and the default e-value of 0.01. For MAGs representing abundant populations and bacteria of functional relevance that we inspected in-depth and manually for sub-cellular profiles of proteins, we automatically annotated the MAGs using the RAST server (98) with default settings with “classic mode”. We choose the following guilds of microorganisms as “functionally relevant” due to their contributions to nutrient removal processes, based on the following rationale and information from the MiDAS Field Guide (38): polyphosphate-accumulating organisms (PAOs) are critical for phosphorus removal (99); nitrifiers and denitrifiers are critical for nitrogen removal (100); and glycogen-accumulating organisms (GAOs) are important because they directly can compete with PAOs for substrates (101). Filamentous bacteria were also included as functionally relevant because they are critical for floc formation and structure (102), and/or problematic “bulking” in activated sludge (103). We defined MAGs that represent “abundant” populations as those with >0.1 average abundance across 23 Danish WTTPs based on Illumina sequence coverage among the 581 non-redundant MAGs, which was performed previously (37) (Supp. Table 1).

### Ortholog group analyses

For orthogroup analyses, protein subsets were subject to OrthoFinder (v 2.3.12) (104) using default settings and “DIAMOND” for the sequence similarity search steps.

### Prediction and annotation of carbohydrate active enzymes

Carbohydrate-active enzymes (CAZymes) were identified using dbCAN 2.0 webserver with HMM, HotPep and DIAMOND detection methods. CAZymes were accepted if hits were obtained by 2 or more of the detection methods. We excluded CAZyme results with probable biosynthetic functions: “Glycoside transferases”, “Soluble_lytic_murein_transglycosylase”, “Peptidoglycan-N-acetylmuramic_acid_deacetylase_PdaC”, “Peptidoglycan_hydrolase_FlgJ”, “Membrane-bound_lytic_murein_transglycosylase_A” and “Membrane-bound_lytic_murein_transglycosylase_D”.

### Prediction and annotation of peptidases

Peptidase/proteases were identified by DIAMOND-BLAST analysis of proteins against the MEROPS database “pepunit_3.lib” (105) with an e-value of 10^-20^. To discern peptidases potentially used for ‘nutrient’ acquisition from other functional roles, e.g., biosynthetic or house-keeping functions, we subjected all predicted peptidases to eggNOG-mapper (as described above) to map them to clusters of orthologous groups (COG). This identified peptidases most similar to catabolic peptidases known for nutrient acquisition, i.e., ‘COG E’ (‘Amino acid transport and metabolism’). We then excluded peptidases that mapped to other categories. We also further removed proteins annotated as “Glutathione hydrolase” that likely have housekeeping functions.

### Prediction and annotation of nucleases and nucleotidases

Nucleases were identified by an iterative approach. First, DIAMOND-BLAST analysis of proteins was performed against a custom seed database (“Nuclease_seed_database.fasta”) (doi.org/10.6084/m9.figshare.25238380) with protein sequences from a previously published study regarding nucleases (70), with an e-value of 10^-10^. Protein sequences of hits were then retrieved and subject to Conserved Domain search tool of CDD to identify and retrieve proteins with similarity to nuclease functional domains, i.e., “endonuclease”, “SNase”, “NUC1”, “SNC”, “HNHC”, “5_nucleotid_C/MPP_superfamily”, “nadN superfamily”, and “PRK09419 superfamily”. Proteins without nuclease domains were discarded. The collected proteins were then added to the DIAMOND-BLAST database, and the proteins were again subjected to DIAMOND-BLAST and Conserved Domain searches to identify additional nuclease proteins. To search for RNase sequences, i.e., iterations of DIAMOND-BLAST analysis of proteins against a custom database (“RNase_seed_database.fasta”) (doi.org/10.6084/m9.figshare.25238380), with an e-value of 10^-10^, collection of hits, and screening for Conserved Domains using the search tool of CDD. Proteins were collected with hits to domains “microbial_RNases superfamily”, “RNase_H_like superfamily”, “RNase_HI_prokaryote_like”, “rnhA”, “RNase_Sa” and “Ribonuclease”. To search for periplasmic nucleotidase-related sequences, i.e., iterations of DIAMOND-BLAST analysis of proteins against a custom database (“PP_nucleotidases_seed_database.fasta”) (doi.org/10.6084/m9.figshare.25238380), with an e-value of 10^-10^, collection of hits, and screening for Conserved Domains using the search tool of CDD. Proteins were collected with hits to domains “MPP_superfamily superfamily”, “5_nucleotid_C”, “ushA”, “MPP_UshA_N_like”, “nadN superfamily” and “PRK09419 superfamily”.

### Prediction and annotation of lipases

Lipases were detected using DIAMOND-BLAST searches of proteins against a custom database (“Lipase_seed_database.fasta”) (doi.org/10.6084/m9.figshare.25238380) based on the ESTHER database (106) and previous work (107), with an e-value of 10^-5^. We also included proteins annotated by Prokka as “Multifunctional_esterase”, “lipase”, and “Glycerophosphodiester_phosphodiesterase”. All proteins with significant hits to potential lipase proteins were subject to Conserved Domain search tool of CDD to identify and retrieve proteins with lipase functional domains, i.e., “EstA”, “Lipase_3”, “SGNH_hydrolase superfamily”, “Abhydrolase”, “GDPD_ScGlpQ1_like”, “ALP_like”, “nSMase”, “PC_PLC”, “PLA1”, “Triacylglycerol_lipase_like”, and “OMPLA superfamily”. Those with “PhoD” domains were not included as lipases.

### Prediction and annotation of heme-binding proteins

To identify predicted secreted cytochromes and other potential heme-binding proteins, we retrieved all proteins predicted to be extra-cytoplasmic locations, as well as those with unknown locations with signal peptides (UNK+SP proteins), and that had “CxxCH” amino acid sequences of typical heme-binding sites (where “x” can be any amino acid). These were retrieved and subject to eggNOG-mapper and the Conserved Domain search tool of CDD, using default setting for both (as described above). Proteins were classified as cytochromes if i) they were annotated as “cytochrome” by Prokka (see above), ii) the eggNOG-mapper functional descriptor contained “cytochrome”, “respiration” and/or other descriptors related to respiration (e.g., denitrification), and/or iii) if they contained cytochrome-type domains as determined by CDD searches with domains including “Cytochrom”, “nanowire_3heme”, “decahem”, “PSCyt1 superfamily”, ”octaheme_Shew superfamily”, or ”MXAN_0977_Heme2 superfamily”. Additionally, we identified many protein sequences with many heme-binding sequences had the “heat shock protein” as an eggNOG-mapper functional descriptor, and therefore they were also collected considering most, but not all, had Conserved Domain hits to cytochrome-like domains.

## Acknowledgements

The project was funded by the Villum Foundation (Dark Matter grant 13351) and Aalborg University.

## Author contributions

Kenneth Wasmund, Conceptualization, Data curation, Investigation, Visualization, Writing – original draft, Writing – review and editing |

Caitlin Singleton, Data curation, Writing – review and editing |

Morten Kam Dahl Dueholm, Data curation, Writing – review and editing |

Michael Wagner, Supervision, Writing – review and editing. |

Per Halkjær Nielsen, Conceptualization, Supervision, Writing – original draft, Writing – review and editing.

## Supplemental Material

Supplement Tables 1-14.

Supp. Fig. 1,2,3,4,5.

## Additional files

### Figshare

Supplement data file 1 - Protein results.xlsx: https://doi.org/10.6084/m9.figshare.25238242

Supplement data file 2 - eggNOG data files - v2.xlsx: https://doi.org/10.6084/m9.figshare.25238248

Protein sequence files = protein sequences from each predicted secreted subcellular compartment: https://doi.org/10.6084/m9.figshare.25238362

Seed protein databases: https://doi.org/10.6084/m9.figshare.25238380

