## Supplementary Figure 1 for "The predicted secreted proteome of activated sludge microorganisms indicate distinct nutrient niches"

**Supp. Fig. 1**. Relationships of counts of proteins from different predicted subcellular locations per MAG with MAG sizes. Each dot represents a MAG.  A) extracellular, B) unknown with signal peptides, C) outermembrane and cell wall, D) periplasmic, E) cytoplasmic membrane, F) cytoplasmic.

**A)**

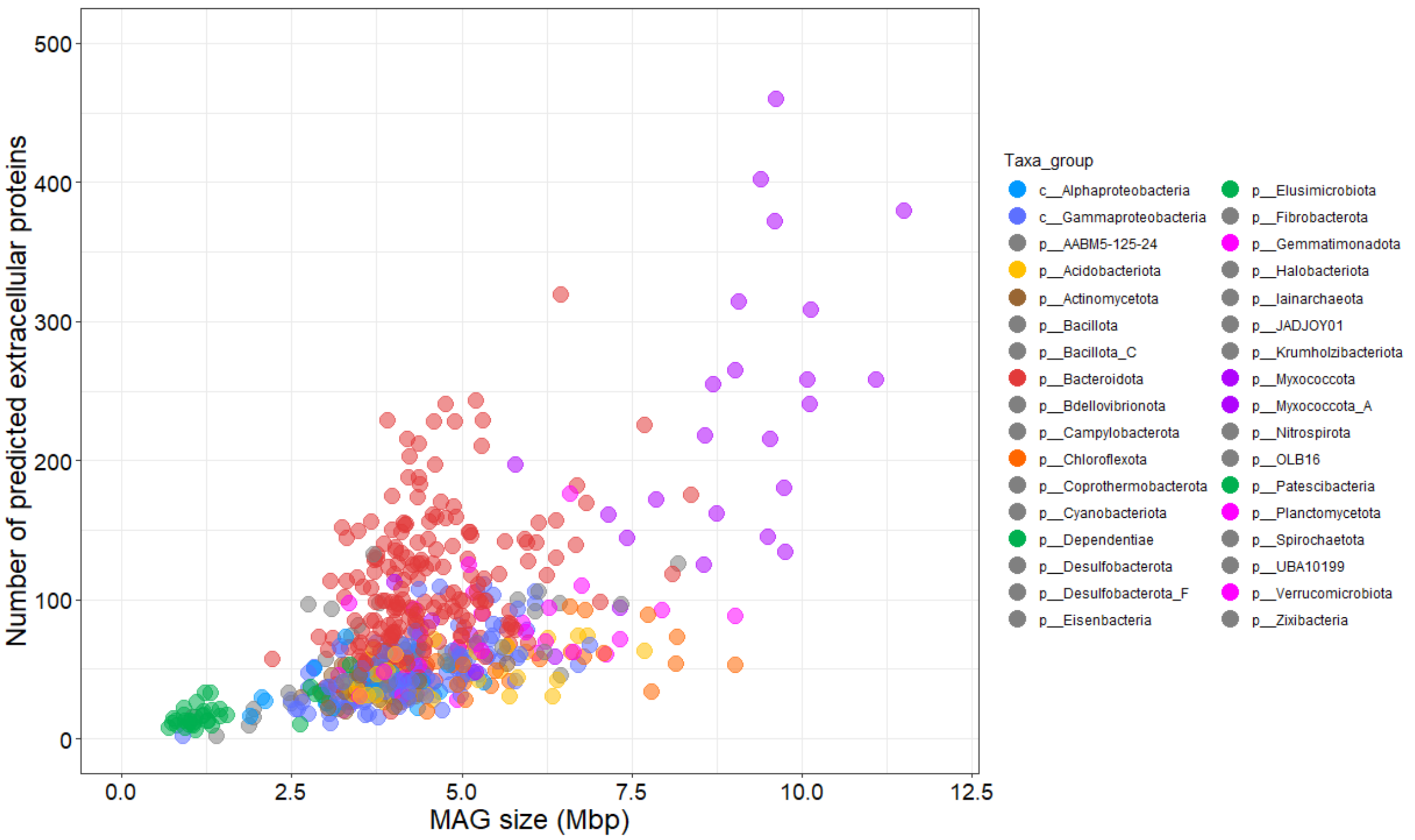

Number of predicted extracellular proteins
MAG size (Mbp)
0
100
200
300
400
500
0.0
2.5
5.0
7.5
10.0
12.5
Taxa_group
c__Alphaproteobacteria
c__Gammaproteobacteria
p__AABM5-125-24
p__Acidobacteriota
p__Actinomycetota
p__Bacillota
p__Bacillota_C
p__Bacteroidota
p__Bdellovibrionota
p__Campylobacterota
p__Chloroflexota
p__Coprothermobacterota
p__Cyanobacteriota
p__Dependentiae
p__Desulfobacterota
p__Desulfobacterota_F
p__Eisenbacteria
p__Elusimicrobiota
p__Fibrobacterota
p__Gemmatimonadota
p__Halobacteriota
p__Iainarchaeota
p__JADJOY01
p__Krumholzibacteriota
p__Myxococcota
p__Myxococcota_A
p__Nitrospirota
p__OLB16
p__Patescibacteria
p__Planctomycetota
p__Spirochaetota
p__UBA10199
p__Verrucomicrobiota
p__Zixibacteria


**B)**

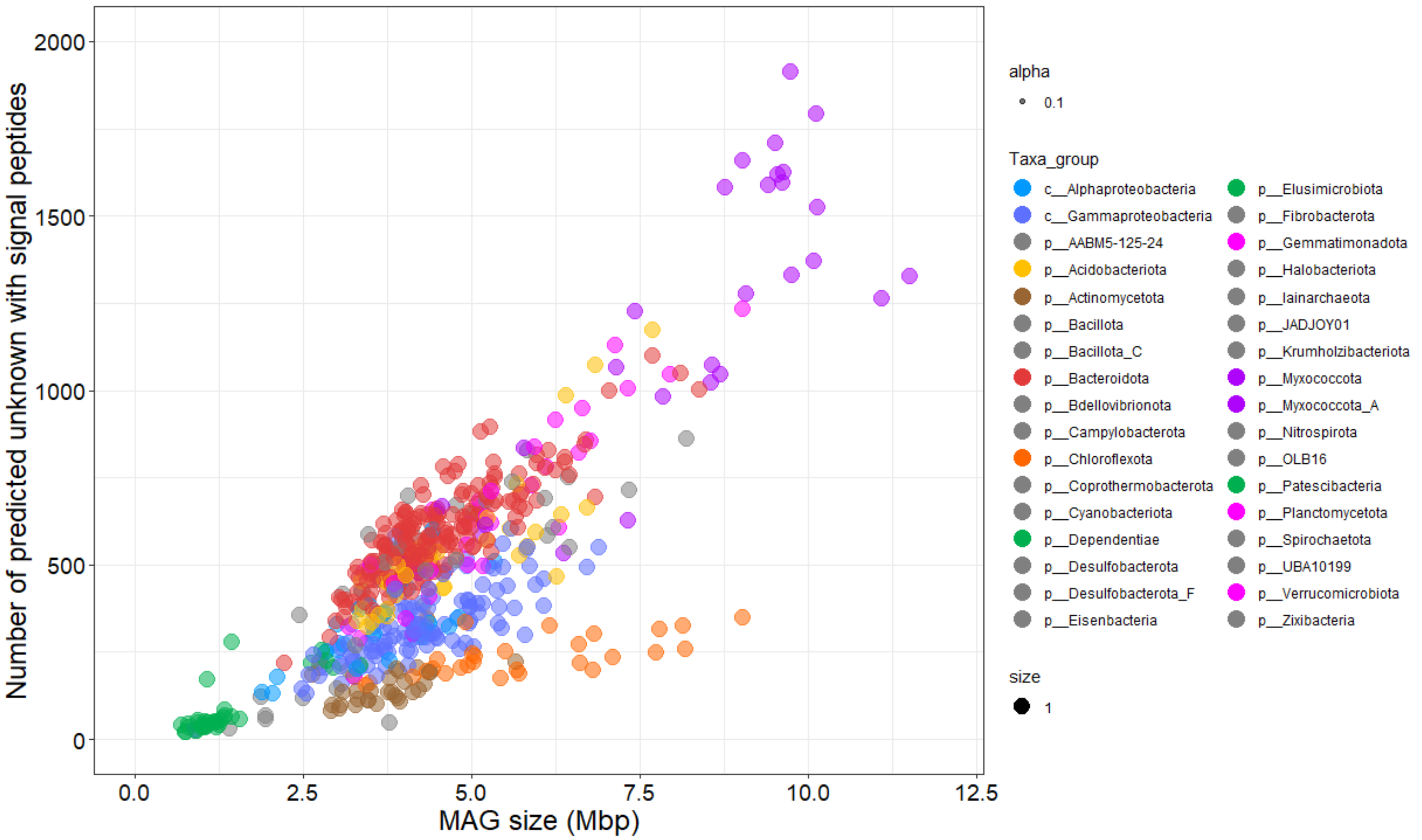

Number of predicted unknown with signal peptides
MAG size (Mbp)
0
500
1000
1500
2000
0.0
2.5
5.0
7.5
10.0
12.5
alpha
0.1
Taxa_group
c__Alphaproteobacteria
c__Gammaproteobacteria
p__AABM5-125-24
p__Acidobacteriota
p__Actinomycetota
p__Bacillota
p__Bacillota_C
p__Bacteroidota
p__Bdellovibrionota
p__Campylobacterota
p__Chloroflexota
p__Coprothermobacterota
p__Cyanobacteriota
p__Dependentiae
p__Desulfobacterota
p__Desulfobacterota_F
p__Eisenbacteria
p__Elusimicrobiota
p__Fibrobacterota
p__Gemmatimonadota
p__Halobacteriota
p__Iainarchaeota
p__JADJOY01
p__Krumholzibacteriota
p__Myxococcota
p__Myxococcota_A
p__Nitrospirota
p__OLB16
p__Patescibacteria
p__Planctomycetota
p__Spirochaetota
p__UBA10199
p__Verrucomicrobiota
p__Zixibacteria
size
1

C)

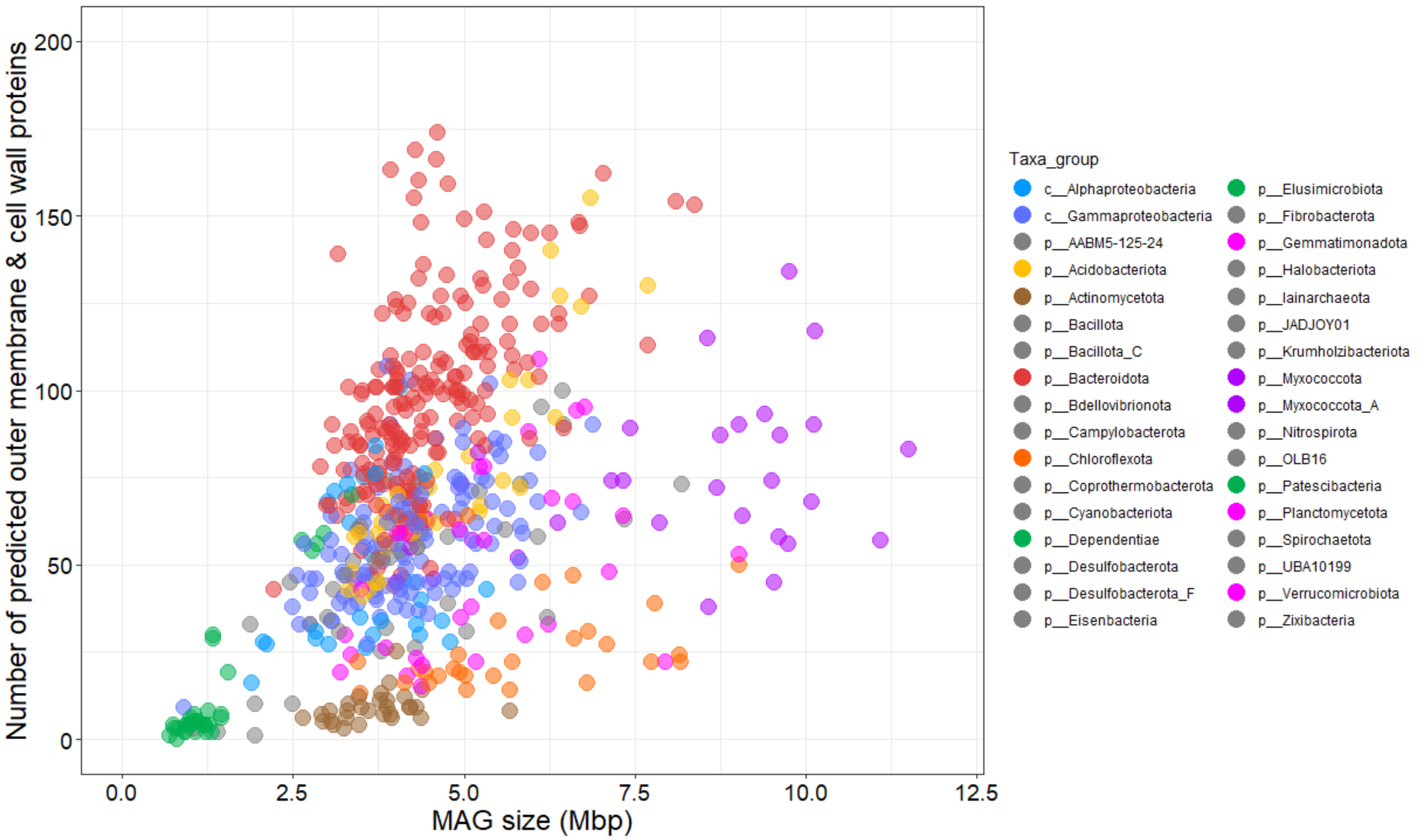

Number of predicted outer membrane & cell wall proteins
200
150
100
50
0
0.0
2.5
5.0
7.5
10.0
12.5
MAG size (Mbp)
Taxa_group
c__Alphaproteobacteria
c__Gammaproteobacteria
p__AABM5-125-24
p__Acidobacteriota
p__Actinomycetota
p__Bacillota
p__Bacillota_C
p__Bacteroidota
p__Bdellovibrionota
p__Campylobacterota
p__Chloroflexota
p__Coprothermobacterota
p__Cyanobacteriota
p__Dependentiae
p__Desulfobacterota
p__Desulfobacterota_F
p__Eisenbacteria
p__Elusimicrobiota
p__Fibrobacterota
p__Gemmatimonadota
p__Halobacteriota
p__Iainarchaeota
p__JADJOY01
p__Krumholzibacteriota
p__Myxococcota
p__Myxococcota_A
p__Nitrospirota
p__OLB16
p__Patescibacteria
p__Planctomycetota
p__Spirochaetota
p__UBA10199
p__Verrucomicrobiota
p__Zixibacteria


D)

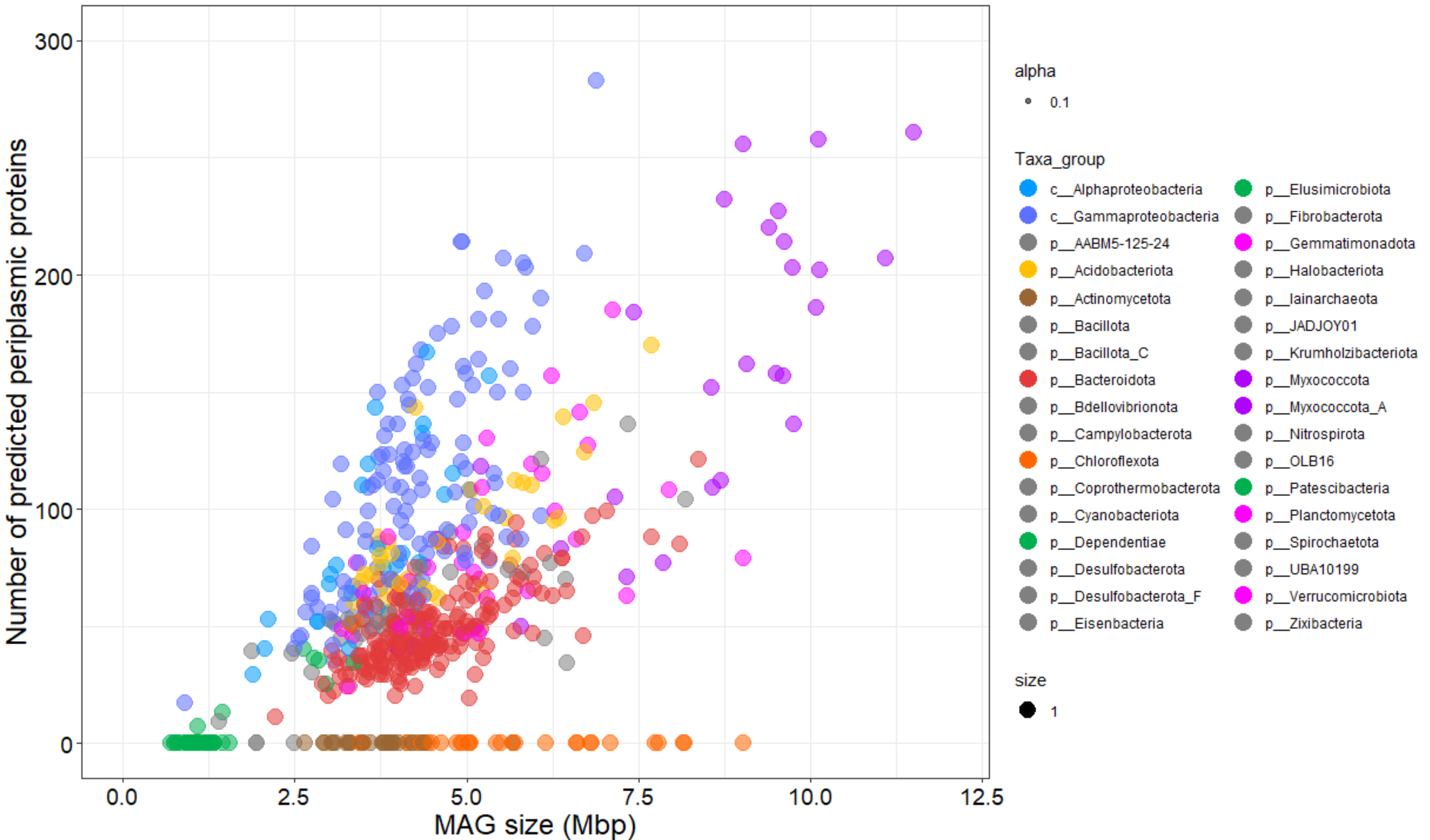

Number of predicted periplasmic proteins
300
200
100
0
0.0
2.5
5.0
7.5
10.0
12.5
MAG size (Mbp)
alpha
0.1
Taxa_group
c__Alphaproteobacteria
c__Gammaproteobacteria
p__AABM5-125-24
p__Acidobacteriota
p__Actinomycetota
p__Bacillota
p__Bacillota_C
p__Bacteroidota
p__Bdellovibrionota
p__Campylobacterota
p__Chloroflexota
p__Coprothermobacterota
p__Cyanobacteriota
p__Dependentiae
p__Desulfobacterota
p__Desulfobacterota_F
p__Eisenbacteria
p__Elusimicrobiota
p__Fibrobacterota
p__Gemmatimonadota
p__Halobacteriota
p__Iainarchaeota
p__JADJOY01
p__Krumholzibacteriota
p__Myxococcota
p__Myxococcota_A
p__Nitrospirota
p__OLB16
p__Patescibacteria
p__Planctomycetota
p__Spirochaetota
p__UBA10199
p__Verrucomicrobiota
p__Zixibacteria
size
1

**E)**

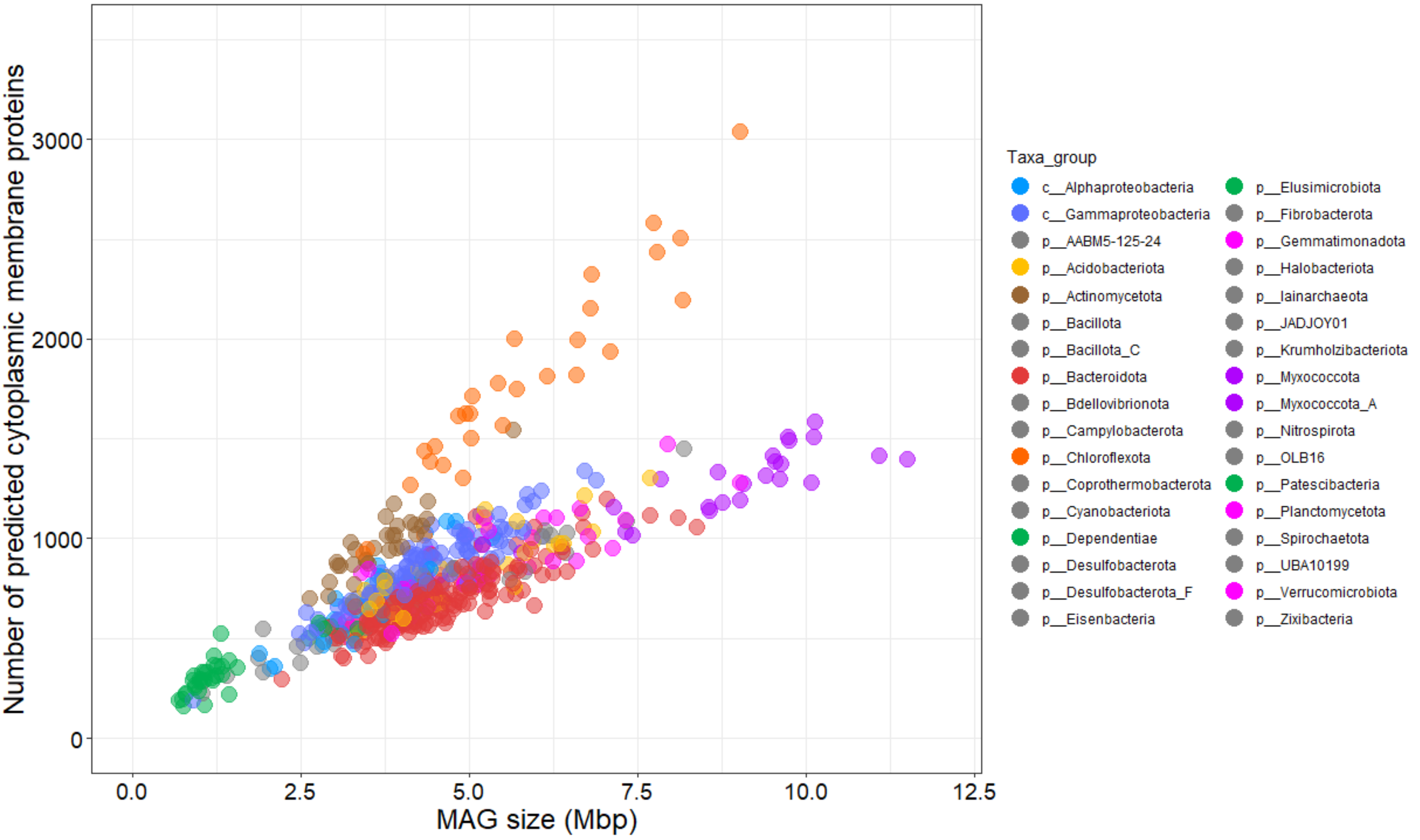
Number of predicted cytoplasmic membrane proteins
0
1000
2000
3000
0.0
2.5
5.0
7.5
10.0
12.5
MAG size (Mbp)
Taxa_group
c__Alphaproteobacteria
c__Gammaproteobacteria
p__AABM5-125-24
p__Acidobacteriota
p__Actinomycetota
p__Bacillota
p__Bacillota_C
p__Bacteroidota
p__Bdellovibrionota
p__Campylobacterota
p__Chloroflexota
p__Coprothermobacterota
p__Cyanobacteriota
p__Dependentiae
p__Desulfobacterota
p__Desulfobacterota_F
p__Eisenbacteria
p__Elusimicrobiota
p__Fibrobacterota
p__Gemmatimonadota
p__Halobacteriota
p__Iainarchaeota
p__JADJOY01
p__Krumholzibacteriota
p__Myxococcota
p__Myxococcota_A
p__Nitrospirota
p__OLB16
p__Patescibacteria
p__Planctomycetota
p__Spirochaetota
p__UBA10199
p__Verrucomicrobiota
p__Zixibacteria

**F)**

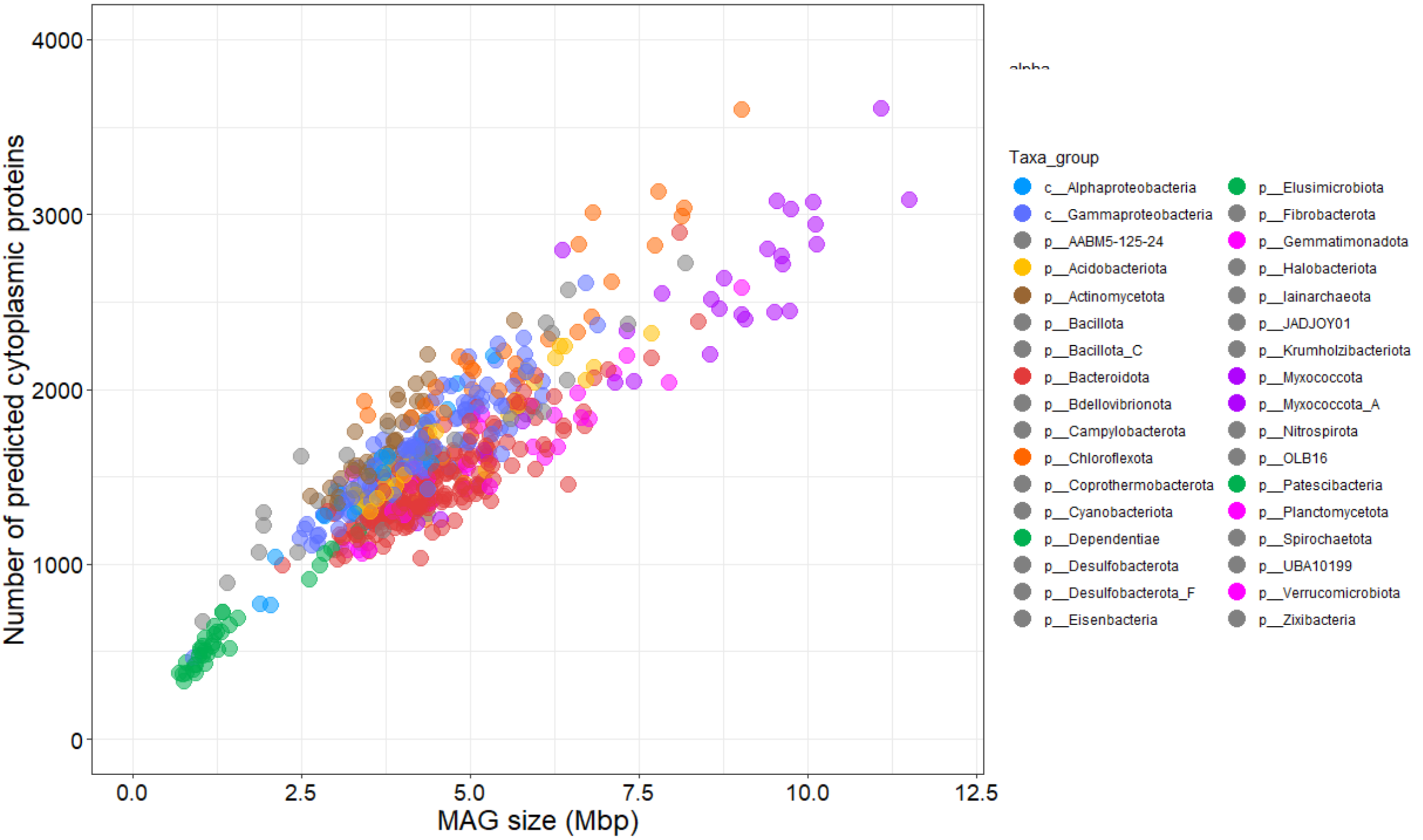
Number of predicted cytoplasmic proteins
0
1000
2000
3000
4000
0.0
2.5
5.0
7.5
10.0
12.5
MAG size (Mbp)
Taxa_group
c__Alphaproteobacteria
c__Gammaproteobacteria
p__AABM5-125-24
p__Acidobacteriota
p__Actinomycetota
p__Bacillota
p__Bacillota_C
p__Bacteroidota
p__Bdellovibrionota
p__Campylobacterota
p__Chloroflexota
p__Coprothermobacterota
p__Cyanobacteriota
p__Dependentiae
p__Desulfobacterota
p__Desulfobacterota_F
p__Eisenbacteria
p__Elusimicrobiota
p__Fibrobacterota
p__Gemmatimonadota
p__Halobacteriota
p__Iainarchaeota
p__JADJOY01
p__Krumholzibacteriota
p__Myxococcota
p__Myxococcota_A
p__Nitrospirota
p__OLB16
p__Patescibacteria
p__Planctomycetota
p__Spirochaetota
p__UBA10199
p__Verrucomicrobiota
p__Zixibacteria
