## Supplementary Figure 2 for "The predicted secreted proteome of activated sludge microorganisms indicate distinct nutrient niches"

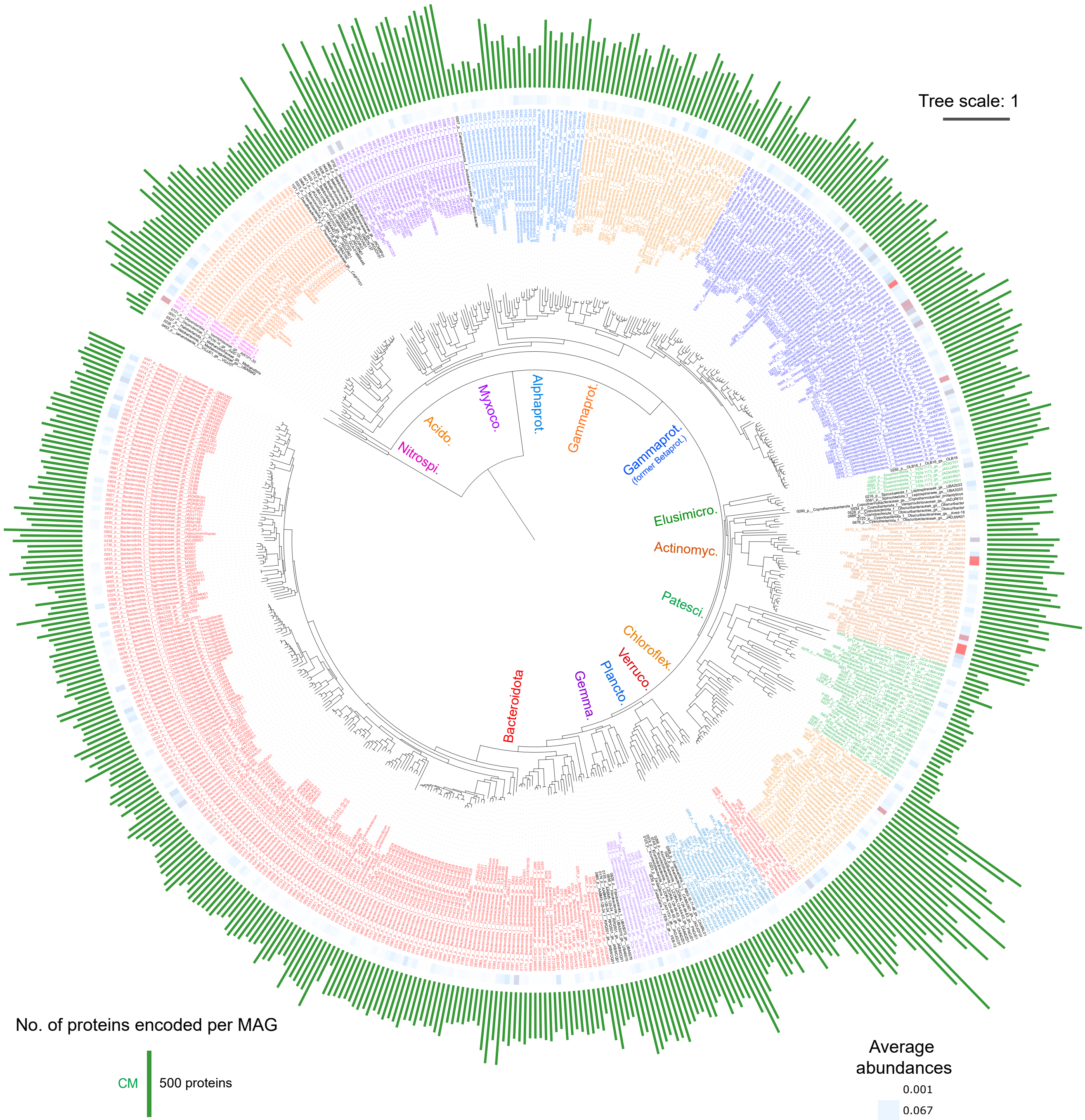


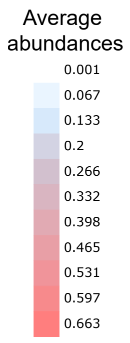


**Supp. Fig. 2**. Phylogenomic tree of 581 MAGs from Danish WWTPs with counts of predicted cytoplasmic memrane proteins. Green bars correspond to counts of predicted cytoplasmic membrane proteins. Most inner rign ("Abund.") with heatmap corresponds to average relative abundances of MAG-populations based on read mapping to MAGs from all metagenomes analysed (values also in Supp. Table 1, colour-scale presented in legend to bottom-right). Leaf labels include the MAG number, followed by taxonomic strings of: phyla (class for Pseudomonadota), family, genus-species, denoted by p__, c__, f__, gs__, respectively. Clades of most major phyla are indicated inside the tree with: Nitrospirota; Acidobact. (Acidobacteriota); Myxococc. (Myxococcota); Alphaprot. (Alphaproteobacteria); Gammaprot. (Gammaproteobacteria); Betaprot. (Betaproteobacteria); Elusimicro. (Elusimicrobiota); Actinomyc. (Actinomycetota); Patesci. (Patescibacteria); Chloroflex. (Chloroflexota), Verruco. (Verrucomicrobiota); Plancto. (Planctomycetota); Gemma. (Gemmatimonadota); Bacteroidota. GTDB species names are only presented if named, i.e., GTDB number codes were removed. The tree is based on a concatenated alignment of protein sequences derived from single copy marker genes obtained from CheckM analysis of MAGs. Scale bar represents 100% sequence divergence.
