## Supplementary Figure 3 for "The predicted secreted proteome of activated sludge microorganisms indicate distinct nutrient niches"

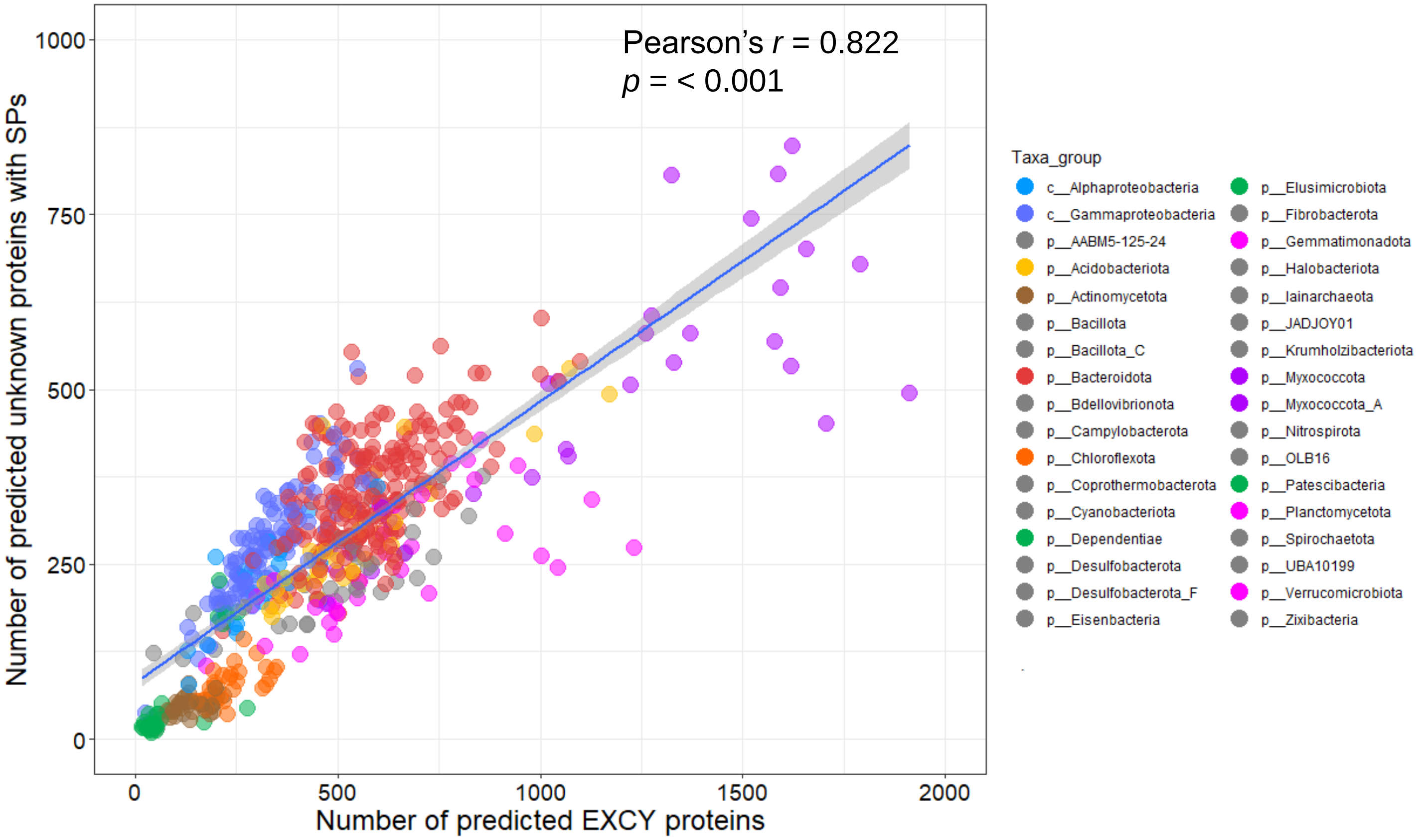


**Supp. Fig. 3**. Correlation of counts of predicted EXCY proteins with those with unknown locations but having signal peptides. "EXCY" = all predicted extracellular, cell wall, outermembrane and periplasmic proteins.
