## Supplementary Figure 4 for "The predicted secreted proteome of activated sludge microorganisms indicate distinct nutrient niches"

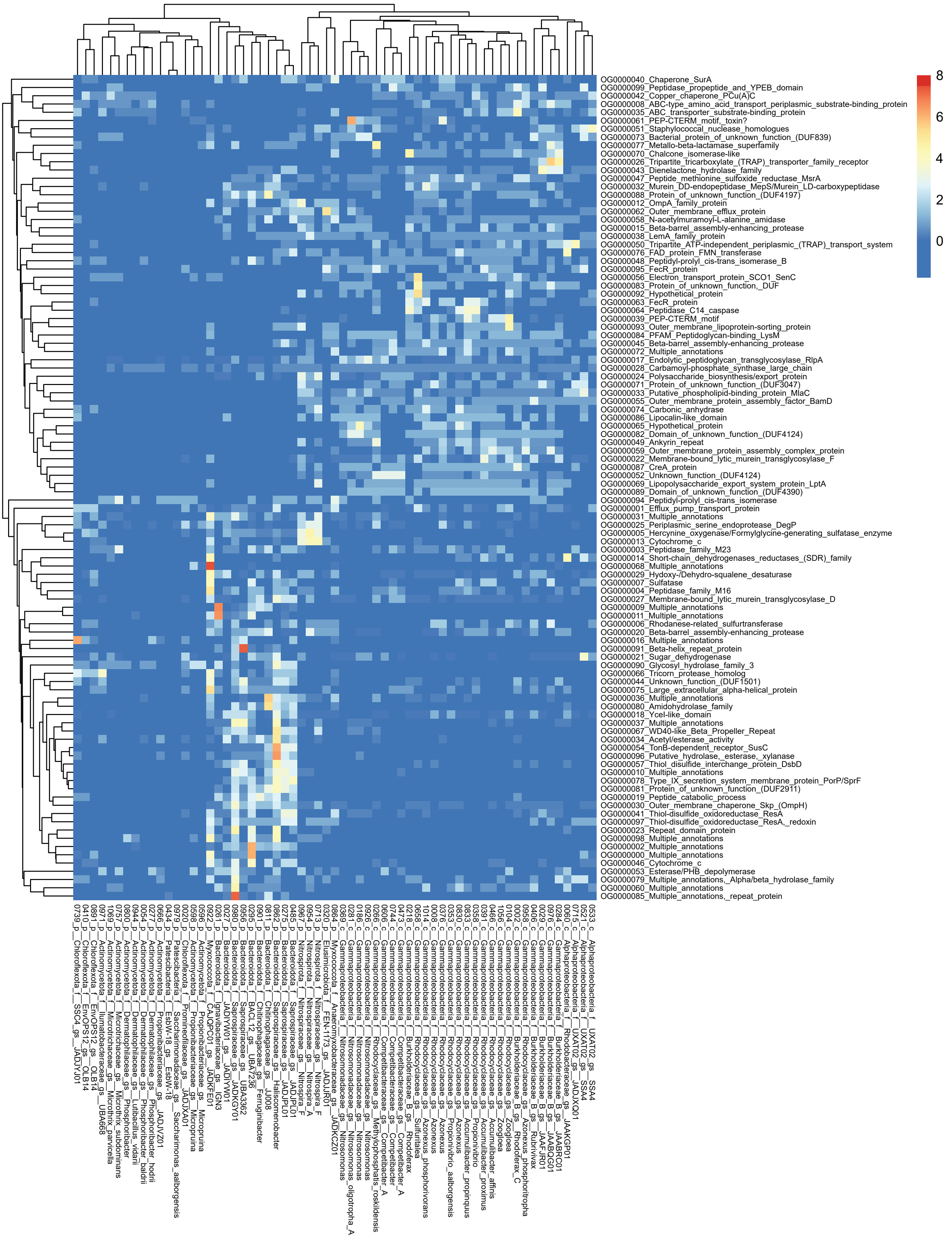


**Supp. Fig. 4**. Heatmap and cluster analysis of proteins from orthogroup analysis of proteins with unknown locations with signal peptides from abundant and functionally relevant taxa. Rows are centered; unit variance scaling is applied to rows. Both rows and columns are clustered using correlation distance and average linkage. The scale for the heatmap colours are indicated in the legend, where the scale maximum of 8 was used to enhance visualisation and differentiation of lower values <8. Note that some values were therefore >8, and raw values of OG counts are available in Supp. Table 3. MAG labels include the MAG number, followed by taxonomic strings of: phyla (class for Pseudomonadota), family, genus-species, denoted by p__, c__, f__, gs__, respectively.
