## SupplementaryInformation for "The predicted secreted proteome of activated sludge microorganisms indicate distinct nutrient niches"

[1] Center for Microbial Communities, Department of Chemistry and Bioscience, Aalborg University, Aalborg, Denmark.
[2] Centre for Microbiology and Environmental Systems Science, Department of Microbiology and Ecosystem Science, University of Vienna, Vienna, Austria.
[3] School of Biological Sciences, University of Portsmouth, Portsmouth, UK.
[#] corresponding.

### Predicted cytoplasmic-membrane-bound proteins

Numbers of predicted cytoplasmic-membrane proteins per MAG were especially high (1504-1937) among various Chloroflexota and Myxococcota MAGs (i.e., those with >2SD above the mean per MAG) (Supp. Table 1 and Supp. Fig. 5). Patescibacteria generally have the fewest per MAG averaging 301±73. The average was 811±342 predicted cytoplasmic-membrane proteins per MAG among all 581 dereplicated MAGs. Ortholog-group (OG) analysis of cytoplasmic-membrane proteins among abundant and "functionally relevant" taxa (*N* = 63 species representatives) (see Methods), identified several highly-represented types of secreted proteins encoded among these MAGs that we suggest could be especially functionally relevant.

A finding of potential interest from the OG analysis of predicted cytoplasmic membrane proteins was that copy numbers of genes for the "low affinity phosphate transporter" (Pit) (OG-71), a potential biomarker for the PAO phenotype (1), are present in higher copy numbers in (≥3 per MAG, and up to 5) (Supp. Table 3) in the most well known PAOs in activated sludge. These include several *Ca*. Phosphoribacter spp. and the *Ca*. Lutibacillus vidarii (MAG 0944) (both former *Tetrasphaera*), and several *Azonexus* (formerly *Ca.* Dechloromonas) and *Ca*. Accumulibacter MAGs. This is noteworthy because Pit is encoded in single copies by many MAGs that don't represent key PAOs, i.e., in 309 of the 581 MAGs of this study. We therefore speculate that having numerous copies of genes for Pit might be a contributing genomic determinant for a PAO phenotype for the respective organisms. This could allow the PAOs to produce extra Pit proteins (2), which could help to increase energy conservation via the proton motive force from the symport of $H^+$ during anaerobic efflux of phosphate (3).

Among other abundant and/or functionally relevant MAGs that are not known PAOs, only a GTDB genus "JAAKGP01" MAG (Alphaproteobacteria) and a *Rhodoferax* MAG (Gammaproteobacteria) also had many copies of *pit* (4 each) (Supp. Table 4). The *Rhodoferax* MAG encodes other key PAO enzymes, i.e., polyphosphate kinase 1 (Ppk1), polyphosphate kinase (Ppk2), and exopolyphosphatase (Ppx), while the JAAKGP01 encodes Ppk1 and Ppk2, but not Ppx. Considering Ppk enzymes are reversible, both MAGs could represent populations with a potential PAO phenotype, although both have low abundances based on average read coverage (Supp. Table 1).

The cluster analysis of OGs and MAGs also revealed several OGs that tended to co-occur with the Pit OG-71 among MAGs (Supp. Fig. 5). Most closely associated was OG-21, which included proteins predicted to be transporter/efflux pumps for $Mg^{2+}/Co^{2+}$. This is intriguing because Pit is a metal-phosphate-dependent co-transporter reliant on divalent cations like $Mg^{2+}$, $Ca^{2+}$, $Co^{2+}$ or $Mn^{2+}$ (4). We therefore speculate the apparent co-dependence for harbouring genes for $Mg^{2+}/Co^{2+}$ transporter may be a mechanism for PAOs in activated sludge to balance the flux of such ions. Genes for proteins of OG-21 also tended to be present in most copies among well known PAOs (Supp. Table 4).

The OG analysis showed that two comammox *Nitrospira* MAGs 0967 and 0713 stood-out for having the most copies of subunits for NADH:quinone oxidoreductase (Nuo) complexes (Supp. Table 4). MAG 0967 has 3 copies of most subunits and one more gene cluster of many Nuo subunits disrupted by a contig break, while MAG 0713 has 3 gene clusters encoding most subunits. Multiple copies have been observed in other comammox genomes (5) and suggest they impart some physiological flexibility to the organisms.

One of the most common proteins among cytoplasmic-membrane OGs were subunits of ABC transporters (Supp. Table 4). When all ABC transporter-related proteins among the OGs were summed for each, the Alphaproteobacteria MAG 0060 had the most ($N$ = 183), followed by various Chloroflexota, Gammaproteobacteria and Actinomycetota MAGs. Numerous ABC-transporter proteins likely impart capabilities to transport diverse organic compounds, as described in the main text.

Finally, we noted many cytoplasmic-membrane OGs related to signalling and sensing, i.e., histidine kinases, and diguanylate and adenylate cyclase sensor proteins (Supp. Table 4). When proteins of these two types were summed, MAGs with many histidine kinase proteins (>1SD above the mean) belonged to Chloroflexota, Bacteroidota, Myxococcota, Gammaproteobacteria and Nitrospirota. For cyclase sensor proteins, MAGs with many (>1SD above the mean) belonged to Gammaproteobacteria. We suggest the differing numbers of these proteins gives indications for numbers or types of external signals these bacteria could respond to.

### Key features of secreted proteomes of the most abundant taxa and key functional groups

Ortholog-group (OG) analysis of extra-cytoplasmic proteins among abundant and "functionally relevant" taxa (*N* = 63 species representatives) revealed the following aspects.

Three of the top 100 OGs contained CAZyme-defined glycoside hydrolases (OG-52 -53 and -60) (Fig. 3 and Supp. Table 2), all of which belonged to GH families with diverse substrate specificities, i.e., GH families 16, 13 and 3. This suggests an importance of secreted glycoside hydrolases for carbohydrate catabolism among many of the abundant members of the communities. MAGs with multiple (≥2) GHs from these OGs mostly belonged to Bacteroidota, Actinomycetota and Chloroflexota (Fig. 3 and Supp. Table 2).

Two abundant OGs had potential catabolic extracellular peptidases, i.e., OG-12 contained S8-family serine peptidases (subtilase family), and OG-48 contained M28 family peptidases (Fig. 3 and Supp. Table 2). Both were common and present among diverse taxa, i.e., among 37 of the 64 MAGs analyzed. These provide an indication for a widespread ability to digest proteins/peptides among abundant populations.

Proteins of the 5'-nucleotidase family with predicted periplasmic or cell wall locations were represented by abundant OG-30 that was spread among various taxa, especially MAGs of Burkholderiales (Gammaproteobacteria), Actinomycetota, and Bacteroidota (Fig. 3 and Supp. Table 2). This suggests the ability to digest nucleotides and/or nucleotide-containing molecules is somewhat common. Nucleotides represent a source of phosphorus, nitrogen and carbon, and released nucleobases and ribose could be salvaged and/or catabolized.

OG-43 contained acyl-homoserine lactone (AHL)-acylase and/or penicillin-acylase related proteins, which were present among diverse MAGs (Fig. 3 and Supp. Table 2). These enzymes may play a role in the quenching of quorum-sensing by hydrolysing AHLs from the environment (6). AHLs play a key role in regulating granule formation in aerobic sludge (7), and thus, microorganisms that degrade AHLs could play an important role in regulating granule and floc formations. Alternatively, some of these enzymes could be used to degrade beta-lactams, as many have shown cross-specificity for both beta-lactams and AHLs (8).

Also relevant for interspecies competition, OG-35 was represented by various proteins with rearrangement hotspot (RHS) repeats, which are typically associated with toxin proteins used for contact-dependent interspecies competition in bacteria (9, 10). These were present in 24 of the 64 MAGs analyzed here, suggesting they have potential to play a role in the lifestyles of many abundant taxa, notably including 3 MAGs of the key nitrogen-cycling *Nitrospira* (Fig. 3 and Supp. Table 2). The

Anaerolineae MAG 0739 (Chloroflexota) had the most copies with 13, suggesting they are especially important for this taxon.

Two abundant OGs contained cytochromes that may facilitate respiration with oxygen, i.e., OG-75 had Cbb3-type cytochrome c oxidase subunit proteins and OG-20 had cytochrome C4 proteins. Both types were present in diverse MAGs (Fig. 3 and Supp. Table 2), although they were both notably absent from most Bacteroidota, present in only 4 of 46 Bacteroidota MAGs. Cytochrome C4 proteins are typically high redox cytochromes and were mainly restricted to Gammaproteobacteria MAGs. Gammaproteobacteria were recently shown to be among the most transcriptionally active organisms in aerobic conditions in activated sludge (11), suggesting these different dominant groups may play different roles under the different phases of redox conditions.

Various OGs with proteins with similarity to T9SS-related proteins were most abundant among Bacteroidota MAGs and clustered together (Fig. 3). T9SS are thought to be distinct to Bacteroidota, and our results therefore suggest they are important for Bacteroidota in WWTP, and may have functions such as secreting hydrolytic enzymes, displaying proteins on cell surfaces (e.g., adhesions), or gliding motility (12). Considering Bacteroidota are abundant and encode diverse secreted proteins, T9SS machinery might be an important determinant for secreting enzymes of Bacteroidota.

### Myxococcota MAGs encode especially large complements of secreted proteins with unknown functions

Myxococcota MAGs encode the highest numbers of predicted extracellular proteins among all MAGs (Fig. 2 and Supp. Table 1). We therefore aimed to briefly explore their predicted extracellular proteins (apart from hydrolytic enzymes that are described elsewhere). We analysed the extracellular protein sequences ($N$ = 5956) from 20 Myxococcota MAGs with the highest predicted extracellular protein complements of any Myxococcota MAGs (from all 1083 MAGs), which range from 197-461 per MAG (Supp. Table 10). To get an overview of potential functions, we subjected the sequences to eggNOG and COG categories (using eggNOG-mapper). Of the 39.5% of extracellular proteins that mapped to COG categories, only few (40.6%) were assigned to functional COG categories (Supp. Table 11). This indicates a very high proportion of the extracellular proteins have no similarity to proteins with known functions. To obtain hints about their properties, we further subjected all top 20 Myxococcota extracellular protein sequences to conserved domain database (CDD) searches. This identified proteins with “Stig1 domains” as the most commonly encoded among the extracellular proteins of the Myxococcota (324 of 5960 proteins analysed) (Supp. Table 12 and Supp. Table 13). Many ($N$ = 171) of these proteins had multiple (≥2) Stig1 domains, with one having as many as 17. Domains of the “myxo_disulf_rpt superfamily” (Myxococcus cysteine-rich repeat) were also commonly detected among the protein set ($N$

= 232) (Supp. Table 12). Both Stigma specific proteins and Myxococcus cysteine-rich repeats are cysteine-rich proteins, and their functions are unknown in bacteria.

### Manual inspections of secreted protein annotations of abundant taxa and key functional groups

In the following section, we performed manual inspections of the predicted secreted proteins of each MAG from the abundant and "functionally relevant" taxa (*N* = 64 species representatives).

**Polyphosphate-accumulating organisms (PAOs)**

PAOs belonging to *Ca*. Phosphoribacter (MAGs 0054 and 0277) and *Ca*. Lutobacillus (MAG 0944) (all formerly "*Tetrasphaera*") (13) represent highly abundant populations critical for P-removal in EPBR systems worldwide (13, 14). Analyses of their predicted extra-cytoplasmic proteins indicate capabilities for catabolic activity for various macromolecules, including proteins, RNA, lipids and a few carbohydrates (Supp. Table 1). All three had genes for extracellular RNases and adjacently encoded cognate cytoplasmic RNase inhibitor proteins ('barnases'), as well as cell wall-bound nucleotidases at other genomic loci. This suggests the capability to break down RNA. Multiple extracellular peptidases may help supply the peptides/amino acids that they are known to ferment (13). All three MAGs encoded single extracellular CAZymes with similarity to alpha-amylases, fitting with previous work that showed starch can be hydrolysed *in situ* by related organisms (15). Predicted cell wall-bound phospholipases (PhoC-type) are also encoded in *Ca*. Phosphoribacter MAGs 0054 and 0277, suggesting a potential to degrade phospholipids.

Secreted polyhydroxyalkanoate (PHA) depolymerases were predicted for all three *Ca*. Accumulibacter MAGs (Supp. Table 14). These PAOs could therefore be capable of digesting extracellular polyhydroxyalkanoates (PHA) (16), which we hypothesize could help them use PHA released from lysed cells, and/or from other taxa. This function has not been shown for these organisms.

PAOs of *Azonexus* (*Ca*. Dechloromonas) have predicted extracellular and/or outer-membrane lipases among all MAGs of this genus (Supp. Table 1). These could help supply fatty acids for beta-oxidation, which was recently predicted for *Azonexus* MAGs (17). The *Azonexus phosphoritropha* (MAG 0958) also had a predicted extracellular PHA depolymerase with an extracellular signal peptide (Supp. Table 14).

**Glycogen-accumulating organisms (GAOs)**

Glycogen-accumulating organisms (GAOs) can compete with PAOs for substrates (e.g., volatile fatty acids), making them undesirable for nutrient removal. MAGs of gammaproteobacterial GAOs including *Ca*. Propionivibrio (MAG 0353 represents the most abundant population) and family *Competibacteraceae* MAGs had generally low numbers of extracellular proteins predicted (Supp. Table 1). GAOs of the *Defluviicoccus* (Alphaproteobacteria) (MAGs 0521, 0533 and 00715) are generally in low abundance, and had few predicted secreted proteins (35-58 per MAG). These proteobacterial GAOs therefore are likely reliant on low molecular weight substrates and/or other VFAs as previously shown (1, 18).

Members of the *Defluviicoccus* (GTDB genus SSA4) (Alphaproteobacteria) are GAOs that are thought to have limited substrate ranges, mainly using VFAs (19). The three MAGs representing this group had relatively low numbers of predicted extracellular proteins (58-35 per MAG), many of which were either flagella related or hypotheticals. PHA depolymerases with TAT signal peptides were identified in all three MAGs (Supp. Table 14). MAG 0521 had a putative extracellular lipase, an extracellular glycerophosphoryl diester phosphodiesterase, and outer-membrane predicted long-chain fatty acid transport proteins that might support use of lipids by this bacterium. Only single predicted extracellular glycoside hydrolases, i.e., alpha-amylase, was found in MAG 0533. These analyses suggest limited substrate ranges for macromolecules for these bacteria.

*Micropruina* (Actinomycetota) are GAOs that can ferment simple sugars and amino acids to drive glycogen storage, and are therefore thought to compete with PAOs of *Ca*. Phosphoribacter and *Ca*. Lutibacillus (former *Tetrasphaera*) that have similar substrate use patterns (20). MAGs of *Micropruina* had a number of predicted hydrolytic exoenzymes, including different nucleases for DNA or RNA, peptidases and CAZymes (Supp. Table 1). Predicted cell wall-bound 5'-nucleotidases were also encoded (Supp. Table 1), and may assist the digestion of nucleic acids. These *Micropruina* MAGs therefore have similar secreted hydrolase profiles to *Ca*. Phosphoribacter and *Ca*. Lutibacillus. This therefore suggests they may also compete with the *Ca*. Phosphoribacter and *Ca*. Lutibacillus PAOs for macromolecules, not just simple sugars and amino acids.

**Nitrogen-dissimilating bacteria**

Nitrogen removal is a key function of WWTPs. Important functional guilds include ‘nitrifiers’ (ammonia- and/or nitrite-oxidizers, including complete ammonia-oxidizers, i.e., ‘comammox’), as well as denitrifiers. Ammonia-oxidizing bacteria (AOB) of the *Nitrosomonas* are typically the most important ammonia-oxidizers in WWTPs. We predicted periplasmic RNases in all four *Nitrosomonas* MAGs (Supp.

Table 14), and we hypothesize they may help them salvage nucleobases, since known members of this group have limited to no heterotrophic capacities (21).

MAGs representing taxa that can contribute to denitrification (*Rhodocyclaceae* of the Gammaproteobacteria) (22, 23), all had predicted extracellular and/or outer-membrane lipases, but few others secreted proteins (Supp. Table 1). Like other Gammaproteobacteria described above, they also generally had many predicted periplasmic proteins, including various substrate-binding proteins of transport systems and enzymes for transforming small molecules like nucleotidases, aromatics and glycerol-3-phosphate, suggesting a capacity to take-up and use small molecules. The two gammaproteobacterial *Zoogloea* MAGs and the JAABQG01 MAG had predicted secreted PHA depolymerases (Supp. Table 14).

**Filamentous bacteria**

Filamentous bacteria are important because: i) they often form a 'backbone' for floc structures; and ii) when in especially high abundances, cause problematic 'bulking' or 'foaming', leading to reduced sedimentation of activated sludge flocs. Our analyses of MAGs of *Ca*. Microthrix (Actinomycetota) identified that apart from secreted lipases described above, these two MAGs and most among the genus, encoded predicted extracellular nucleases (Supp. Table 1). While lipolytic lifestyles are known for these organisms (24, 25), nucleic acid-degrading capabilities are not known. Filamentous members of the Anaerolineae (Chloroflexota) have numerous predicted secreted glycoside hydrolases, as well as extracellular or cell wall nucleases and nucleotidases, and numerous peptidases (although not all were classified as "nutrient-acquiring") (Supp. Table 1), suggesting versatile catabolic capacities for macromolecules. Potential for anaerobic polysaccharide utilization is consistent with previous MAR-FISH studies showing this function for Anaerolineae (26). Analysis of three *Haliscomenobacter* spp. (Bacteroidota) MAGs strongly suggested they are polysaccharide-degraders, with MAG 0485 having the second most predicted extracellular CAZymes of any MAG ($N$ = 39), while the three MAGs also had high numbers of predicted outer-membrane and periplasmic CAZymes. Together, these findings suggest many of the key filamentous bacteria in activated sludge have capacity to be primary-degraders of organic macromolecules.

Seven additional MAGs representing abundant Bacteroidota were analyzed (MAGs 0027, 0261, 0295, 0811, 0901, 0956 and 0980). In general, these MAGs were among those that encoded the highest numbers of predicted extracellular and outer-membrane proteins among all MAGs (Supp. Table 1), with many encoding various TonB-receptor proteins and/or porins, indicating high capacities to translocate diverse and possibly large molecules.

Examination of protein types most enriched from OGs revealed Ignavibacterium MAG 2061 and Chitinophagales MAG 0295 had expansive numbers of predicted secreted proteins with T9SS sorting domains (43 and 37, respectively), suggesting type-9 secretion systems maybe important for their ecological interactions.

Several MAGs were analysed representing abundant filamentous Anaerolineae (Chloroflexota), i.e., MAG 0739 (*Ca*. Amarolinea), MAGs 0410 and 0891 (*Ca*. Villigracilis/GTDB genus 'OLB14'), and MAG 0020 (Unc. Promineofilaceae). The *Ca*. Amarolinea MAG 0739 had relatively large predicted secreted protein complements, with 93 predicted extracellular proteins. These four Chloroflexota MAGs all encoded multiple secreted glycoside hydrolases, as well as extracellular or cell wall nucleases and nucleotidases, and numerous peptidases (although not COG-E), suggesting versatile hydrolytic capacities for macromolecules (Supp. Table 1). Potential for anaerobic polysaccharide utilization is consistent with previous MAR-FISH studies showing this function (27).

***Abundant Bacteroidota***

Seven MAGs representing abundant Bacteroidota were analysed here, with five MAGs from uncharacterised Chitinophagales (MAGs 0027, 0295, 0811, 0956 and 0980), one *Ferruginibacter* MAG 0901 (order Chitinophagales), and one *Ignavibacterium* MAG 0261 (class Ignavibacteria). All had numerous predicted extracellular proteins (ranging from 53 to 104 per MAG), with MAGs 0295, 0980, 0261 having among the highest predicted extracellular protein complements (between 104, 99 and 73, respectively) in the whole dataset (Supp. Table 1). They also had among the highest numbers of outer-membrane proteins predicted (93-163 per MAG), with only MAGs 0027 and 0901 having fewer (67 each). Many were annotated as TonB-receptor proteins and/or porins, indicating high capacities to translocate diverse and possibly large molecules. These abundant Bacteroidota encoded numerous CAZymes that probably target various polysaccharides including some from plants, as described above. Notable exceptions were MAG 0027 (uncharacterised family of Chitinophagales) and MAG 0956 (family Saprospiraceae), which have few predicted secreted CAZymes, respectively (Supp. Table 1). Both had multiple predicted secreted peptidases (COG-E), and both had extracellular nucleases (Supp. Table 1).

Notably, the abundant Saprospiraceae MAG 0980 encoded a cytochrome with >100 heme-binding sites (Supp. Table 8). These abundant Bacteroidota MAGs also had numerous copies of cytochrome-c551 peroxidases, likely used for detoxification of peroxides. *Ignavibacterium* MAG 0260 had numerous predicted extracellular and periplasmic multi-heme cytochromes, and could help explain a previous observation for their enrichment when iron(III) was added to anaerobic wastewater (28).

***Abundant taxa - other***

In addition to gammaproteobacterial PAOs and GAOs described above, a number of other MAGs represent abundant Gammaproteobacteria, belonging to families *Rhodocyclaceae* and *Burkholderiaceae* (both former "Betaproteobacteria") (MAGs 0029, 0266, 0406, 0658 and 0976). Common features of these MAGs included relatively low numbers of extracellular proteins (18-58 per MAG), low numbers of outer-membrane proteins (33-71 per MAG), and high numbers of periplasmic proteins (84-156 per MAG) (Supp. Table 1). Few secreted hydrolases for macromolecules were predicted, with only 2 MAGs having predicted outer-membrane lipases. The MAGs generally encode numerous and different types of predicted periplasmic proteins/subunits with probable substrate-binding activities, e.g., components of TRAP- and ABC-transporter complexes. These were often annotated (automatically) to have specificities for peptides and amino acids. Together with limited or absent extracellular hydrolases, it suggests these abundant Gammaproteobacteria are specialized to take-up relatively small and freely available molecules.

Further, we analyzed three MAGs of abundant populations from recently described phyla, i.e., Elusimicrobiota (MAG 0320) and Patescibacteria (MAGs 0979 and 0434). These MAGs have very few predicted extracellular proteins (9-13 per MAG), with only the two Patescibacteria MAGs each encoding predicted extracellular nucleases (Supp. Table 1). They therefore have limited capacity to process larger organic molecules outside of their cells. The Elusimicrobiota MAG had relatively many predicted outer-membrane proteins ($N$ = 57) and porins/barrel like proteins compared to its small genome (2.62 Mbp) and relative to Gammaproteobacteria, for example (average genome size = 4.26 Mbp, and average 57 outer-membrane proteins). This possibly suggests they can nevertheless import or transport various molecules.

Two Myxococcota MAGs represent abundant populations, i.e., *Anaeromyxobacter* MAG 0864 (midas_s_1192) and MAG 0922 (midas_s_814) of uncharacterised GTDB order UBA796. Compared to many of the other Myxococcota, *Anaeromyxobacter* MAG 0864 had few predicted extracellular proteins ($N$ = 48), although it has numerous predicted outer-membrane and periplasmic proteins (82 and 118, respectively), with relatively many outer-membrane porins/channels ($N$ = 13) (Supp. Table 1). *Anaeromyxobacter* MAG 0864 encodes a few extracellular and UNK+SP CAZymes, together at two genomic loci with other periplasmic and cytoplasmic glycoside hydrolases and carbohydrate transporters, suggesting co-ordinated capabilities to digest some polysaccharides. The *Anaeromyxobacter* MAG 0864 also had the second most extracellular cytochrome proteins of any MAG ($N$ = 8), and also numerous predicted periplasmic cytochromes ($N$ = 32). This indicates high redox flexibility, as discussed previously for the cultured relative *A. dehalogenans* (29). Notably, no reductive dehalogenase homologs were identified that could enable respiration of halogenated organics, and that are present in *A. dehalogenans* (29).

In comparison, Myxococcota MAG 0922 has the largest array of predicted extracellular proteins (*N* = 161) among the most abundant/core organisms analysed here. It has extensive extracellular hydrolytic potential, encoding three predicted extracellular peptidases, few extracellular and UNK+SP CAZymes, as well as potential lipases and nucleases among UNK+SP proteins (Supp. Table 1). OG and protein domain analyses revealed a large array of predicted extracellular proteins (*N* = 57) with probable adhesion properties, as well as RTX toxin-like proteins (*N* = 27) (Supp. Table 2).

Actinomycetota MAG 0971 represents an abundant and core population in Danish WWTPs (midas_s_1015) (14), belonging to an undescribed genus of the Ilumatobacteraceae. The MAG had a relatively low number of predicted extracellular proteins (41) and cell wall proteins (6) (Supp. Table 1). It was distinct from other abundant Actinomycetota, in that it encoded no secreted lipases or CAZymes, but a single extracellular nuclease was encoded. Most other extracellular proteins were hypotheticals making predictions difficult. The OG analysis identified multiple AHL acylases/beta lactamases.
